# Detection of uncoupled circadian rhythms in individual cells of *Lemna minor* using a dual-color bioluminescence monitoring system

**DOI:** 10.1101/2020.06.10.143727

**Authors:** Emiri Watanabe, Minako Isoda, Tomoaki Muranaka, Shogo Ito, Tokitaka Oyama

## Abstract

The plant circadian oscillation system is based on the circadian clock of individual cells and coordinates the circadian behavior of the plant body. To observe the cellular circadian behavior of both the oscillator and its output in plants, we developed the dual-color bioluminescence monitoring system that automatically measured the luminescence of two luciferase reporters simultaneously at a single-cell level. We selected a yellow-green-emitting firefly luciferase (LUC+) and a red-emitting luciferase (PtRLUC) that is a mutant form of Brazilian click beetle ELUC. We used *AtCCA1::LUC*+ and *CaMV35S::PtRLUC* to observe the cellular behavior of the oscillator and output, respectively. These bioluminescent reporters were introduced into the cells of a duckweed, *Lemna minor*, by particle bombardment. Time series of the bioluminescence of individual cells in a frond were obtained using a dual-color bioluminescence monitoring system with a green-pass- and red-pass filter. Luminescence intensities from the LUC+ and PtRLUC of each cell were calculated from the filtered luminescence intensities. We succeeded in reconstructing the bioluminescence behaviors of *AtCCA1::LUC*+ and *CaMV35S::PtRLUC* in the same cells. Under prolonged constant light conditions, *AtCCA1::LUC*+ showed a robust circadian rhythm in individual cells in an asynchronous state in the frond, as previously reported in studies using other plants. In contrast, *CaMV35S::PtRLUC* stochastically showed circadian rhythms in a synchronous state. Thus, we clearly demonstrated the uncoupling between the oscillator and output in individual cells. This dual-color bioluminescence monitoring system is a powerful tool to analyze various stochastic phenomena accompanying large cell-to-cell variation in gene expression.

**Significance statement:** We succeeded in establishing the world’s first dual-color bioluminescence monitoring system at a single-cell level that enables simultaneous measurement of the luminescence activities of two reporter genes in plants. This system is a strong tool to analyze stochastic phenomena, and we clearly demonstrated the uncoupling of rhythmic behavior between two bioluminescent reporters in individual cells that stochastically occurred in the same plant.

## Introduction

Many physiological phenomena in plants show daily rhythms under day/night cycles, and they show circadian rhythms even under constant conditions. The circadian clock is composed of a number of clock genes that form transcription–translation feedback loops in individual cells (Nagel and Kay, 2012). Thus, the circadian clock is basically a cell-autonomous system. These concepts have been led by various studies using bioluminescence monitoring techniques (Muranaka and Oyama, 2018). Bioluminescence rhythms of transgenic plants expressing a firefly luciferase under the control of circadian promoters are observed using a photomultiplier tube or a high-sensitive camera. Bioluminescence imaging of transgenic plants can spatially distinguish circadian rhythms in a plant body or a specific organ; damping (spontaneous desynchronization) of bioluminescence rhythms in a region and asynchronous rhythms in an organ have been clearly shown (Fukuda *et al*., 2007; Fukuda *et al*., 2012; Wenden *et al*., 2012). These observations suggested heterogeneous circadian behavior among cells even in the same tissue. The heterogeneity was quantitatively demonstrated by the observation of bioluminescence circadian rhythms at a single-cell level in duckweed plants (Muranaka *et al*., 2013; Muranaka and Oyama, 2016; Okada *et al*., 2017; Isoda and Oyama, 2018). A bioluminescent reporter, *AtCCA1::LUC*+, was used for gene transfection to cells with the particle bombardment method. The *Arabidopsis CCA1* gene is one of the clock genes and is highly expressed around dawn (Wang and Tobin, 1998). *CCA1* is widely conserved among plants and its function in circadian rhythm is also conserved (Miwa *et al*., 2006; Serikawa *et al*., 2008; McClung, 2013; Linde *et al*., 2017). This reporter activity, which shows a robust circadian rhythm with a high amplitude (Nakamichi *et al*., 2004; Muranaka *et al*., 2015), was monitored for circadian rhythms of individual cells by using an automated imaging system with an EM-CCD camera (Muranaka *et al*., 2013; Muranaka and Oyama, 2020). Duckweeds are suitable for imaging the entire plant because of their tiny, flat, floating bodies. In addition, duckweed fronds (leaf-like structure) are immobile in the vertical direction, making it unnecessary to adjust the focus during long-term monitoring (Hillman, 1961; Muranaka *et al*., 2013). Among duckweed species, *Lemna gibba* has long been used for studies on circadian rhythms (Muranaka and Oyama, 2018). Recently, *Lemna minor*, which is phylogenetically close to *L. gibba*, has been used for various studies because an efficient genetic transformation system has been established and the genome sequence information is available (Chhabra *et al*., 2011; Van Hoeck *et al*., 2015).

As a basis of the clock regulation of plant physiology, a relatively large number of genes are regulated by the circadian clock (Covington *et al*., 2008; Michael *et al*., 2008). Among these clock-regulated genes, large variations in peak phases, amplitudes, and waveforms are observed. They are likely to be regulated by other signals, such as light, temperature, and developmental events; the gene expression of many are altered in tissues, developmental stages, and environmental conditions. Variation in the circadian behavior of gene expression among genes has been studied in detail by monitoring transgenic plants carrying a circadian bioluminescent reporter (Thain *et al*., 2000; Hall *et al*., 2002; Michael and McClung, 2002; Thain *et al*., 2002; Muranaka and Oyama, 2018). Although the variation or heterogeneity in the circadian behavior occurs in individual cells, comparative analysis among circadian-regulated genes has not been performed at the single-cell level. Analysis of the correlation of circadian traits (phase, period, stability, etc.) between various circadian behaviors at the single-cell level is necessary to reveal the cell-autonomous basis of the circadian system. Especially, cells that are asynchronous in a tissue in a plant are an interesting target for analysis. A dual-color bioluminescence monitoring system is a tool to simultaneously detect the expression levels of two reporter genes; the circadian activities of different promoters of cyanobacteria and mice have been analyzed using this system (Kitayama *et al*., 2004; Nakajima *et al*., 2004, 2005; Ogura *et al*., 2005; Kwon *et al*., 2010; Nishide *et al*., 2018). Simultaneous detection of the cellular circadian rhythms of two bioluminescent reporters was only reported for mammalian cultured cells (Kwon *et al*., 2010). Kwon *et al*. developed a dual-path optical luminescence imaging system and simultaneously tracked two circadian gene expression patterns for several days in individual cells. They used two luciferases: a green-emitting ELuc derived from *Pyrearinus termitilluminans*, and a red-emitting SLR derived from *Phrixothrix hirtus*. The bioluminescences of single cells that were transfected with two mammalian clock-gene reporters showing different phases were separated into green- and red-color images using dichroic mirrors and imaged on the CCD sensor in the microscope system.

In this study, we developed a dual-color bioluminescence monitoring system that enabled us to approach different circadian traits in individual cells in a plant. Based on the single-cell bioluminescence imaging system that was previously developed by Muranaka and Oyama (2016), we additionally set a filter changer to separate luminescence from different color luciferases in the same sample. We chose the commonly used firefly luciferase (LUC+) and the red-shift mutant (S246H/H347A) of ELuc from *Pyrearinus termitilluminans* (PtRLUC) as the luciferase reporters (Nishiguchi *et al*., 2015). LUC+ and PtRLUC emit yellow-green (λ_max_ = ~560 nm) and red (λ_max_ = 602 nm) luminescences, respectively. As a robust circadian bioluminescent reporter, we chose *AtCCA1::LUC*+, which was previously used for the observation of circadian rhythms at a single-cell level in a duckweed plant (Muranaka *et al*., 2013; Muranaka and Oyama, 2016; Okada *et al*., 2017; Isoda and Oyama, 2018). This reporter activity basically represents the behavior of the circadian clock in individual cells; it is regarded as a direct output of the cellular oscillator. As an output reporter with a circadian rhythm, we chose *CaMV35S::PtRLUC* in which PtRLUC was driven by the *CaMV35S* promoter. While the *CaMV35S* promoter is known as a constitutive promoter, the *CaMV35S::LUC*+ reporter showed a bioluminescence circadian rhythm in duckweed plants under constant light after light/dark cycles (Muranaka *et al*., 2015). *CaMV35S* is a virus-derived promoter that is not directly related to the clock oscillation; this reporter activity appears to reflect an output downstream of the clock. By co-transfecting cells in *L. minor* with *AtCCA1::LUC*+ and *CaMV35S::PtRLUC*, we succeeded in monitoring two reporter activities in individual cells of a plant. The results of the dual-color bioluminescence monitoring system clearly indicated the uncoupling of circadian rhythms between *AtCCA1::LUC*+ and *CaMV35S::PtRLUC* in the same cells.

## Results

### Characterization of bioluminescence of cells expressing LUC+ or PtRLUC in *Lemna minor*

We first confirmed that the transfection procedure of particle bombardment for *L. gibba* worked on *L. minor* (Figure S1a). The major target of this transfection was epidermal cells (Figure S1b,c). Approximately 70% of transfected cells in the fronds were epidermal cells and the remainder were mesophyll cells. Then, by using an automatic luminescence monitoring system with photomultiplier tubes, we roughly confirmed that *LUC*+ and *PtRLUC* under the *CaMV35S* promoter showed a circadian rhythm in *L. minor* similarly to that reported for *L. gibba* (Figure S2, Muranaka *et al*., 2015). As reported for *CaMV35S::LUC*+ in *L. gibba, CaMV35S::LUC*+ and *CaMV35S::PtRLUC* in *L. minor* showed circadian rhythms under constant light after light/dark cycles at the whole plant level (Figure S2a,b). In contrast, both reporters in the *L. minor* plants that had not experienced any light/dark cycles showed almost no circadian rhythmicity (Figure S2c,d). Thus, *CaMV35S::LUC*+ and *CaMV35S::PtRLUC* showed similar bioluminescence traces in *L. minor*.

To separately monitor LUC+ and PtRLUC bioluminescence in the same cells, we set up a dual-color bioluminescence monitoring system. As shown in Figure 1a, we set a filter changer to the single-cell bioluminescence imaging system that was previously reported by Muranaka and Oyama (2016). Using this system, we roughly examined the luminescence spectra of *CaMV35S::LUC*+ and *CaMV35S::PtRLUC* (Figure 1b). One day after the transfection, the luminescence intensities of a frond were measured with a series of optical band-pass filters. In plant cells, maximum emission wavelengths were around 560 nm and 600 nm for LUC+ and PtRLUC, respectively. Based on the spectra of luminescence, we selected a set of band-pass filters for LUC+ (green-pass filter: 490–570 nm) and PtRLUC (red-pass filter: 610–650 nm). The foci for a sample were altered when the filters were set. The focus for each filter was optimized by adjusting the distance between the sample and the lens by driving a z-axis motor for the stage (Figure 1a). Figure 1c shows an example of a set of images for an *AtCCA1::LUC*+-transfected frond and a *CaMV35S::PtRLUC*-transfected frond at a time in monitoring. Exposure times for each image were optimized to obtain luminescence within an appropriate range. These sets of images were captured every 1 h and individual luminous spots were quantified for each filtered image. Figure 1d shows a set of time series data for the luminescence intensity of a spot (green arrow in Figure 1c) in the *AtCCA1::LUC*+-transfected plant under constant light conditions (Exp1-1 in Table S1). Intensities of luminescence captured without a filter (hereafter unfiltered luminescence intensities, *L*; gray line) showed a robust circadian rhythm for a week. Intensities of luminescence transmitted through the green-pass filter (hereafter green luminescence intensities, *L_g_*; green line) also showed a robust circadian rhythm with a similar waveform to that of unfiltered luminescence intensities. Intensities of luminescence transmitted through the red-pass filter (hereafter red luminescence intensities, *L_r_*; red line) were much lower than the unfiltered luminescence intensities but showed a circadian rhythm. Figure 1e shows a set of time series data for the luminescence intensity of a spot (red arrow in Figure 1c) in the *CaMV35S::PtRLUC*-transfected plant under constant light conditions (Exp2-1 in Table S1). Unfiltered luminescence intensities (gray line) showed a low-amplitude circadian rhythm with a high-frequency noise. Compared to the unfiltered luminescence intensities, the green luminescence intensities (green line) were close to zero. The red luminescence intensities (red line) showed a circadian rhythm with a similar waveform to that of the unfiltered luminescence intensities.

**Figure 1.**
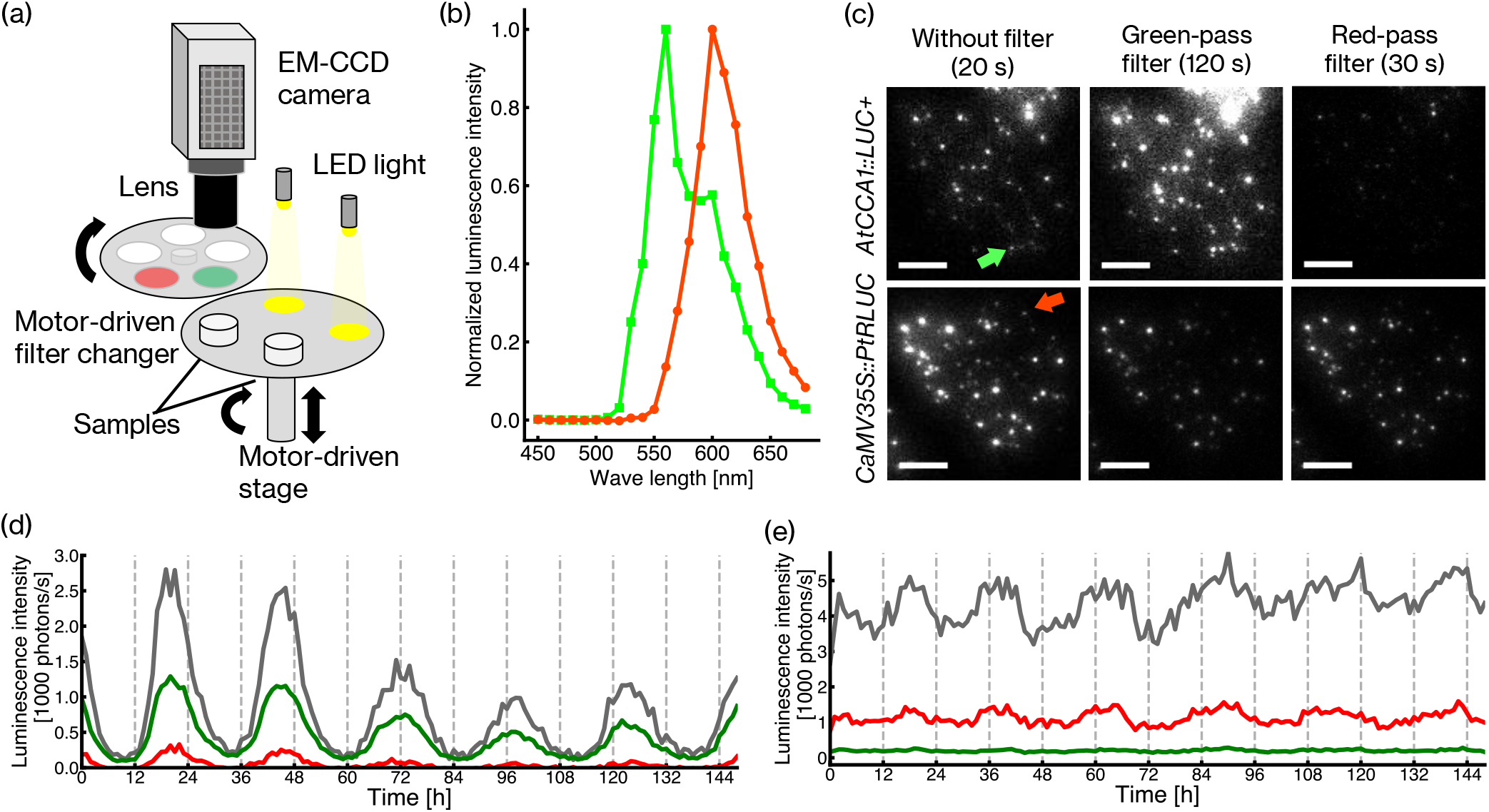
Measurement of bioluminescence of LUC+ and PtRLUC in *Lemna minor*. (a) Schematic drawing of the imaging device for the dual-color bioluminescence monitoring system. Sample dishes containing duckweed plants are set on a motor-driven stage. The stage rotates to change sample positions underneath the lens of an EM-CCD camera or an end of optic fibers guiding LED light. A motor-driven filter changer is set between the sample and the lens. The focus of an image for the sample is adjusted by a z-axis motor. (b) Luminescence spectra of LUC+ and PtRLUC expressed in *L. minor*. (c) Images of luminous spots in a frond of *L. minor* transfected with *AtCCA1::LUC*+ (upper row; Exp1-1) or *CaMV35S::PtRLUC* (lower row; Exp2-1). A luminescence image of a frond without filter (left), and that of the same area filtered with a green-pass filter (490–570 nm) (middle) or red-pass filter (610–650 nm) (right) are shown. The exposure times for each filter condition are represented in parentheses. The green- and red-arrows indicate the luminous spots shown in (d) and (e), respectively. (d, e) Time series data of luminescence intensities of a luminous spot for *AtCCA1::LUC*+ (d) and *CaMV35S::PtRLUC* (e) in each filter condition. Luminescence images for each filter condition were captured every 1 h. Gray, green, and red lines represent the unfiltered, green, and red luminescence intensities, respectively. The illumination conditions for the duckweed plant during luminescence monitoring was constant light except for 18-min hourly darkness for luminescence capturing. Images in (c) were captured at time 0 in (d) and (e). Bars represent 1 mm.

### Calculation to separately monitor LUC+ and PtRLUC bioluminescence in the same cells

First, we quantitively analyzed the transmittance of the filters by using time series data of the *AtCCA1::LUC*+ transfected cells (Exp1-1 in Table S1). The transmittance of a data point for the two filters (*L_g_/L* for the green-pass filter, *L_r_/L* for the red-pass filter) was calculated for those data with *L* > 1000 photons/s. The time series of *AtCCA1::LUC*+ luminescence included data with very low intensities due to its high-amplitude rhythm, and the transmittance of those data points was widely distributed owing to being close to the background level of their unfiltered- and/or filtered luminescence intensities (Figure S3a). Figure 2a shows histograms of the transmittance for the green-pass (green bins) and red-pass (red bins) filters. The transmittance of LUC+ was normally distributed for both filters, with an average of 0.49 ± 0.06 for the green-pass filter (Kolmogorov–Smirnov, *P* = 0.62) and 0.09 ± 0.02 for the red-pass filter (Kolmogorov–Smirnov, *P* = 0.25). The normal distribution suggests that these mean values can be used as the representative values of their transmittance. The scatter plot of green- and red-pass filter transmittance for each data point did not show correlation between them (Figure 2b). Note that the transmittance of LUC+ appeared to be affected by the expression level (Figure S4). The luminescence intensities of *CaMV35S::LUC*+ expressing cells were ~100 times higher than those of *AtCCA1::LUC*+, and the transmittance of *CaMV35S::LUC*+ looked to be lower and higher for the green-pass- and red-pass filter, respectively. Then, we quantitively analyzed the transmittance of the filters by using the time series data of the *CaMV35S::PtRLUC*-transfected cells (Exp2-1 in Table S1). Luminous spots showed strong emissions and did not include those with intensities close to the background level (Figure S3b). The transmittance of PtRLUC was normally distributed for the red-pass filter, with an average of 0.26 ± 0.03 (Kolmogorov–Smirnov, *P* = 0.51) (Figure 2c). The transmittance for the green-pass filter was low, with an average of 0.045 ± 0.007. The Kolmogorov–Smirnov *P*-value for the transmittance of the green-pass filter was very low (*P* = 0.00), suggesting non-normal distribution (Figure 2c). This seems to be due to the difficulty in obtaining the exact values of transmittance for individual spots with very low intensities of green-filtered luminescence. The scatter plot of green- and red-pass filter transmittance for each data point did not show correlation between them (Figure 2d). For these reasons, we used these mean values as the representative values of their transmittance.

**Figure 2.**
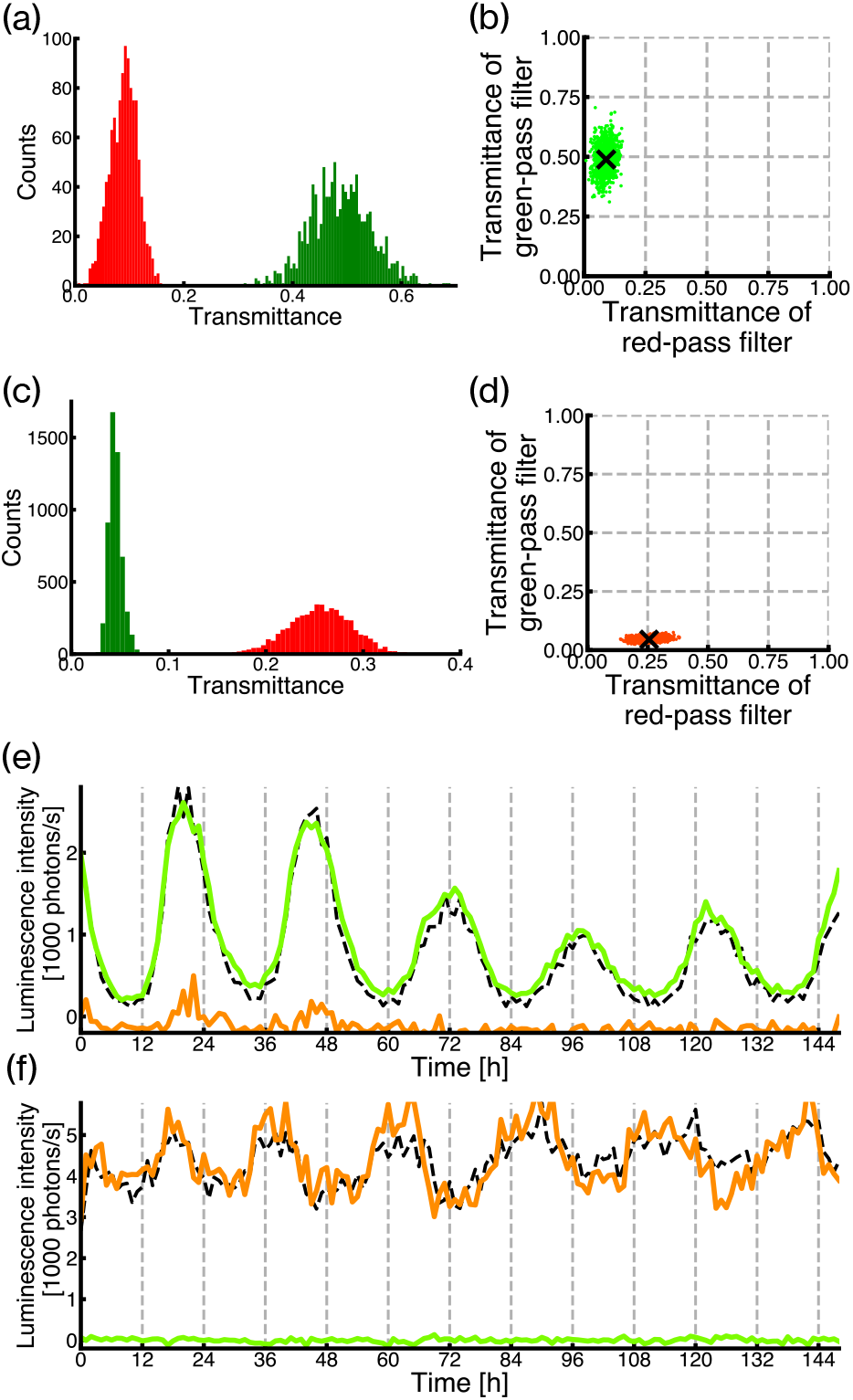
Estimation of luminescence intensities of LUC+ or PtRLUC by using filtered luminescence intensities. Time series data of luminescence intensities obtained in Exp1-1 and Exp2-1 were used. Twenty-eight (LUC+) and 38 (PtRLUC) luminous spots with 149 time-lapse data points (149 h) were calculated for transmittance. Those data including one or more pixels with a saturated signal in the spot area for one or more filter conditions were excluded. (a) Histograms for the transmittance of LUC+ for green-pass (green bars) and red-pass filters (red bars). Data with unfiltered luminescence intensities (*L*) more than 1000 photons/s are counted (total 1129 data points). The bin size is 0.05. (b) A scatter plot of transmittance of LUC+ for the green-pass filter versus that for the red-pass filter. The cross mark shows the average transmittance. (c) Histograms for the transmittance of PtRLUC for the green-pass (green bars) and red-pass filters (red bars). The data condition and the bin size are the same as (b) (total 5354 data points). (d) A scatter plot of transmittance of PtRLUC for the green-pass filter versus that for the red-pass filter. The cross mark shows the average transmittance. (e, f) Reconstruction of time series data of luminescence intensities of a luminous spot for *AtCCA1::LUC*+ (e) and *CaMV35S::PtRLUC* (f). Luminescence of LUC+ (green line) and PtRLUC (red line) were reconstructed using the time series data of filtered luminescence intensities that are represented in Figure 1d,,e. A dashed line in each graph represents the original time series data of unfiltered luminescence intensities.

Next, we provided a calculation formula to estimate the luminescence intensities from two luciferases (*L_LUC+_, L_PtRLUC_*) in the same cells from the filtered luminescence intensities (*L_g_, L_r_*). As explained in Figure S5,

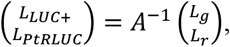

where the elements of matrix A are transmittance for the combinations of luciferases and filters. To confirm that the formula worked well, time series data for estimated luminescence intensities were compared to those of unfiltered luminescence intensities (*L*) for the cells that were separately transfected with *AtCCA1::LUC*+ (Exp1-1 in Table S1) or *CaMV35S::PtRLUC* (Exp2-1 in Table S1). As shown in Figures 2e and S6 on the whole, the *L_LUC+_* values in the time series of *AtCCA1::LUC*+ transfected cells matched with *L*, and the *L_PtRLUC_* values were close to 0. Thus, the formula worked well to reconstruct *AtCCA1::LUC*+ luminescence from the filtered luminescence intensities. However, at time points with strong intensities (*L* > ~5000 photons/s), the *L_LUC+_* and *L_PtRLUC_* values tended to be underestimated and overestimated, respectively (arrows in Figure S6). This appeared to be caused by the expression-level-dependent alteration of LUC+ transmittance (Figure S4). Thus, it would be better to use an appropriate matrix for the calculation depending on the expression level of LUC+. As shown in Figures 2f and S7, the *L_PtRLUC_* values in time series of the *CaMV35S::PtRLUC*-transfected cells were well matched with *L*, and the *L_LUC+_* values were kept close to 0. Thus, the formula worked well to reconstruct *CaMV35S::PtRLUC* luminescence from the filtered luminescence intensities. Therefore, we concluded that the calculation formula was practical for estimating the luminescence intensities of the two reporters.

### Estimation of luminescence intensities of the two reporters in the co-transfected cells

We measured the luminescence of the cells co-transfected with the two reporters, with and without filters in a time series. Figure 3a shows an example of a luminous spot exhibiting a circadian rhythm. The red luminescence intensities were higher than the green ones in most time points. Figure 3b shows a plot of the transmittance of the green- and red-pass filters at every time point for the spot. In the plot, the green- and red crosses show the representative transmittance values of LUC+ and PtRLUC luminescence, respectively (see Figure 2b,d). Points on the line segment connecting these crosses (blue line) satisfy the condition where the sum of *L_LUC+_* and *L_PtRLUC_* is *L* when *L_LUC+_* and *L_PtRLUC_* are reconstructed using the calculation formula. Data points are distributed close to the line segment, suggesting that the calculation formula works to estimate the luminescence intensities of the two reporters in the same cell. Note that many data points are around the endpoint of the line segment (red cross) because the PtRLUC luminescence dominated over the LUC+ luminescence for this cell. Figure 3c shows the time series of reconstructed luminescence intensities of the two reporters for this cell. Both *AtCCA1::LUC*+ and *CaMV35S::PtRLUC* showed circadian rhythms, while the luminescence intensities of *CaMV35S::PtRLUC* were higher than those of *AtCCA1::LUC*+ in most time points. As observed in single-reporter transfection experiments, *CaMV35S::PtRLUC* showed a low-amplitude rhythm with a high-frequency noise and *AtCCA1::LUC*+ showed a high-amplitude rhythm. The *AtCCA1::LUC*+ rhythm appears to include a jaggy noise probably due to the partial interference of *CaMV35S::PtRLUC* luminescence.

**Figure 3.**
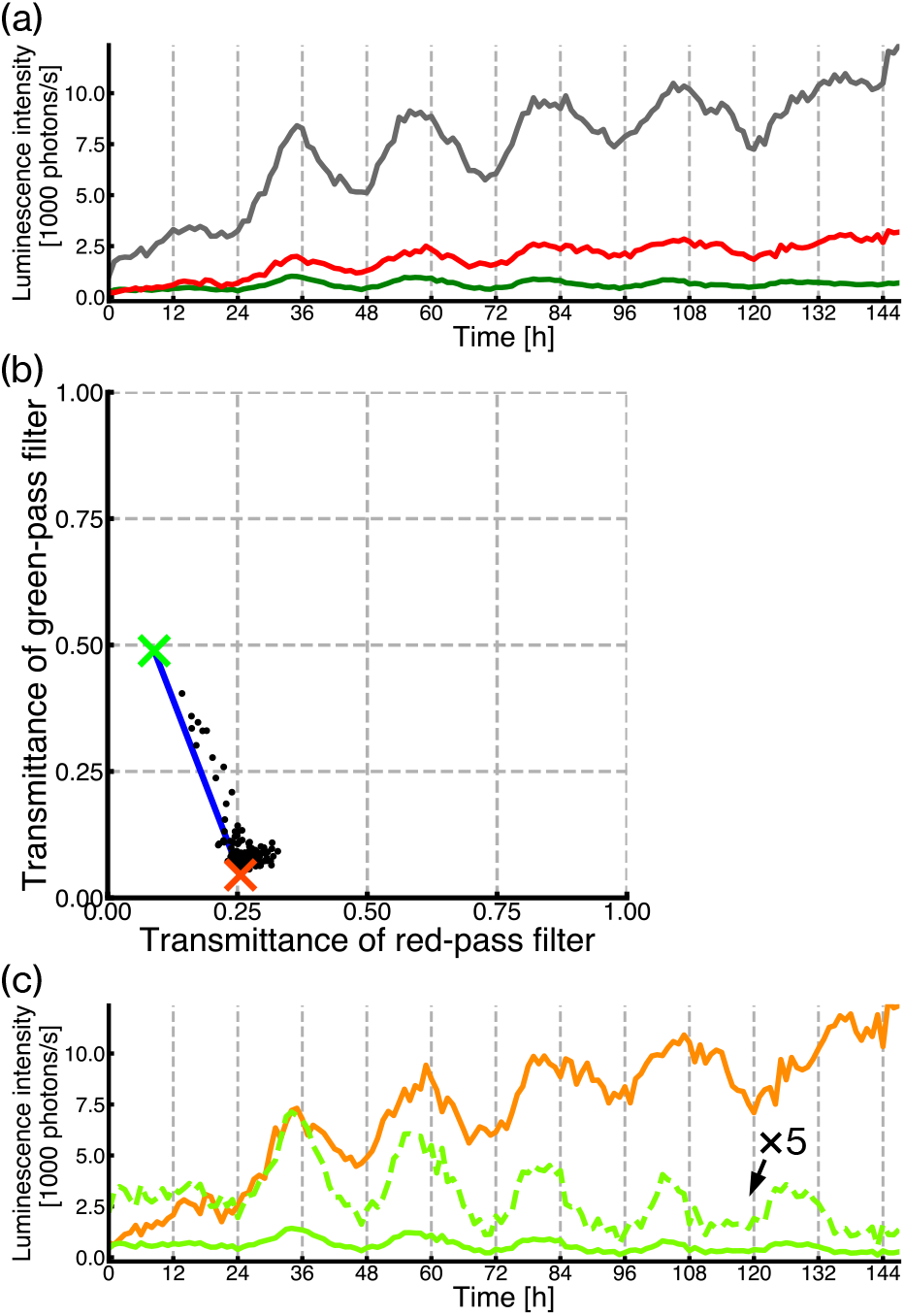
Reconstruction of luminescence intensities for *AtCCA1::LUC*+ and *CaMV35S::PtRLUC* of a co-transfected cell. Images of luminous spots in a frond of *L. minor* co-transfected with *AtCCA1::LUC*+ and *CaMV35S::PtRLUC* (Exp3-1) were captured and analyzed for a luminous spot with 149 time-lapse data (149 h). The growth, illumination, and monitoring conditions were the same as Figure 1d,e. (a) Time series data of luminescence intensities of a luminous cell in each filter condition. Luminescence intensities of the luminous spot are plotted [gray for without filter (*L*), green for green-pass filter (*L_g_*), red for red-pass filter (*L_r_*)]. (b) A scatter plot of transmittance for the green-pass filter versus that for the red-pass filter. Black dots represent the transmittance of individual data. Cross marks show the average transmittance for *AtCCA1::LUC*+ (green, Figure 2b) or *CaMV35S::PtRLUC* (red, Figure 2d) in single-reporter transfection experiments. Dots on the blue line segment satisfy the formula: *L_LUC+_* + *L_PtRLUC_* = *L*. (c) Reconstructed time series data of luminescence intensities for *AtCCA1::LUC*+ (light-green line) and *CaMV35S::PtRLUC* (orange line). A dashed light-green line represents the five-fold magnification of LUC+ luminescence.

Figure S8 shows the time series of reconstructed luminescence intensities of the two reporters for 34 cells in a frond (Exp3-1). For each cell, the overall luminescence was higher in the time series of *CaMV35S::PtRLUC* than in that of *AtCCA1::LUC*+, although the difference varied among cells. Based on the results obtained from the single-reporter transfection experiments, we decided on the following three criteria for luminous spots that were used for time series data analysis of both reconstructed luminescences (Figure S9). (1) Spots with extremely low expression levels of either reporter were removed. (2) Spots showing an overly high expression of LUC+ were removed because of the expression-level-dependent alteration of its transmittance for filters. (3) Spots with luminescence ratios of LUC+ and PtRLUC in an appropriate range were selected. The number of analyzable luminous spots varied among co-transfection experiments; 3 to 20 cells were selected in six independent experiments (Figure S8, Table S1).

### Evaluation of reconstructed bioluminescence traces of the two reporters in co-transfection experiments

We first performed rhythm analysis on bioluminescence traces by following the scheme in Figure S9. The free-running period (FRP) and relative amplitude error (RAE) of each luminescence trace were calculated using the fast Fourier transform non-linear least squares (FFT-NLLS) method (Plautz *et al*., 1997). The RAE value represents the degree of confidence of rhythmicity, ranging from 0 (complete sine-fitting) to 1 (arrhythmic). In this analysis, bioluminescence traces with RAE < 0.2 are defined as rhythmic, and the rest are defined as arrhythmic (Figure S10). In single-reporter transfection experiments on *AtCCA1::LUC*+, bioluminescence traces of most cells showed high-amplitude rhythms (Figure S6), and the proportion of rhythmic traces to analyzable ones ranged from 0.93 to 0.97 in the four independent experiments (Figure 4a). Thus, every cell in *L. minor* basically showed the self-sustained circadian rhythm of this reporter under constant light, as has been reported in *L. gibba* (Muranaka and Oyama, 2016). In contrast, luminous spots of *CaMV35S::PtRLUC* in single-reporter transfection experiments showed traces with low-amplitude circadian rhythms or without circadian rhythmicity (Figure S7), and the proportion of rhythmic traces to analyzable ones ranged from 0.29 to 0.55 in four independent experiments (Figure 4a). The bioluminescence traces of *CaMV35S::PtRLUC* judged as rhythmic (RAE < 0.2) showed lower amplitudes and rougher waveforms than did those of *AtCCA1::LUC*+ (Figure S10). Thus, the circadian rhythm of *CaMV35S::PtRLUC* was unstable and the rhythmicity occurred stochastically in the cells of plants under prolonged constant light conditions. Since the proportion of rhythmic cells for *CaMV35S::PtRLUC* was clearly lower than that for *AtCCA1::LUC*+, it was expected that *CaMV35S::PtRLUC* luminescence traces without circadian rhythmicity would be observed in cells showing the *AtCCA1::LUC*+ rhythm in the co-transfection experiments.

**Figure 4.**
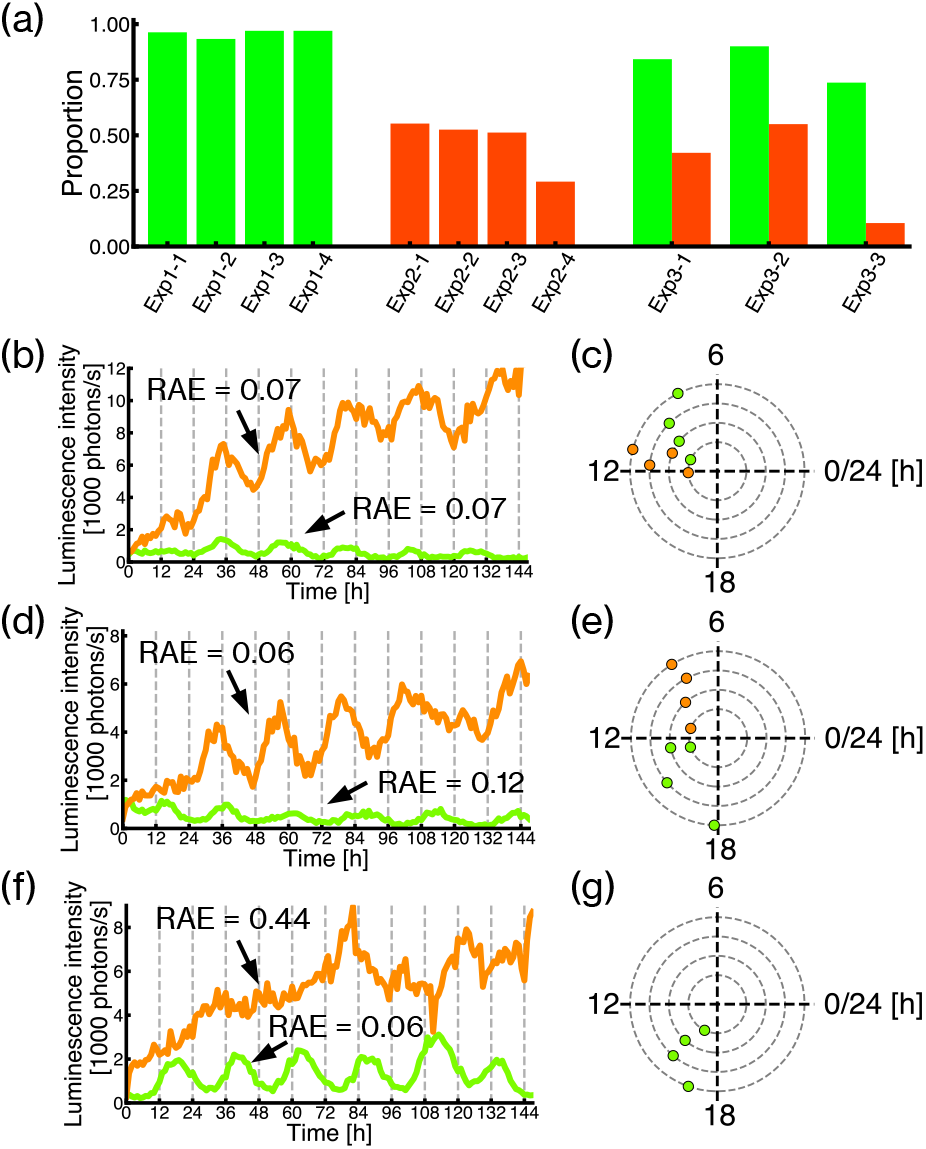
Quantitative analysis of cellular rhythms for *AtCCA1::LUC*+ and *CaMV35S::PtRLUC* of co-transfected cells. (a) Circadian rhythmicity of cellular bioluminescence traces of *AtCCA1::LUC*+ and *CaMV35S::PtRLUC*. Red and green bars represent proportions of cells showing bioluminescence traces with circadian rhythms (RAE < 0.2) in single-reporter transfection experiments for *AtCCA1::LUC*+ (Exp1-1–1-4) and *CaMV35S::PtRLUC* (Exp2-1–2-3), and co-transfection experiments for both reporters (Exp3-1-3-3). (b, d, f) Examples of reconstructed time series data of luminescence intensities for *AtCCA1::LUC*+ (light-green line) and *CaMV35S::PtRLUC* (orange line) in a co-transfection experiment (Exp3-1). The RAE value for each bioluminescence trace is represented. (c, e, g) Time of peaks of individual cellular bioluminescence rhythms of *AtCCA1::LUC*+ (light green) and *CaMV35S::PtRLUC* (orange). The peak time(s) shown in day 2 (24–48 h) is plotted on the innermost circle and those in day 3 (48–72 h), day 4 (72–96 h), and day 5 (96–120 h) are plotted on the second, third, and fourth circles, respectively. The peak times of the bioluminescence rhythms (RAE < 0.2) in (b), (d) and (f) are represented in (c), (e) and (g), respectively.

In six co-transfection experiments on *AtCCA1::LUC*+ and *CaMV35S::PtRLUC*, three experiments had more than 10 analyzable luminous spots for both reconstructed luminescences (Table S1). In these three experiments, the proportion of rhythmic traces of *AtCCA1::LUC*+ to analyzable ones ranged from 0.74 to 0.90, while that of *CaMV35S::PtRLUC* ranged from 0.11 to 0.55 (Figure 4a). The proportion for *AtCCA1::LUC*+ looked lower in co-transfection experiments than in single-reporter transfection ones. There appeared to be interference from *CaMV35S::PtRLUC* luminescence in *AtCCA1::LUC*+ traces to increase their RAE values. In fact, reconstructed *AtCCA1::LUC*+ luminescence traces in co-transfection experiments included jaggy noises, which were often observed in *CaMV35S::PtRLUC* luminescence traces (Figures S6, S7, S8).

As expected in single-reporter transfection experiments, *CaMV35S::PtRLUC* luminescence traces without circadian rhythmicity were observed in cells showing the *AtCCA1::LUC*+ rhythm. As shown in Figures 4b,d,f and S8, some cells showed robust *AtCCA1::LUC*+ rhythms and low-amplitude *CaMV35S::PtRLUC* rhythms, while some showed circadian rhythmicity only in *AtCCA1::LUC*+. Reconstructed *CaMV35S::PtRLUC* luminescence traces in co-transfection experiments appeared to be similar to those observed in single-reporter transfection experiments. In fact, peak times of reconstructed *CaMV35S::PtRLUC* luminescence traces occurred seemingly independent of those of *AtCCA1::LUC*+ rhythms (Figure 4c,e,g). Thus, interference from *AtCCA1::LUC*+ luminescence appeared low enough to maintain the original properties of *CaMV35S::PtRLUC*. Taken together, it is demonstrated that the luminescence behaviors of the two reporters can be simultaneously monitored in individual cells by using the dual-color bioluminescence monitoring system.

### Uncoupling of circadian behavior between *AtCCA1::LUC*+ and *CaMV35S::PtRLUC* in the same cells

While *AtCCA1::LUC*+ reflects the behavior of cellular oscillator, the rhythm of *CaMV35S::PtRLUC* can be regarded as a circadian oscillatory output (Muranaka *et al*., 2015; Muranaka and Oyama, 2016). The presence of both cells with and without the *CaMV35S::PtRLUC* rhythm in the same frond indicates that the cellular output observed in *CaMV35S::PtRLUC* luminescence can be uncoupled from the cellular oscillator to behave arrhythmically. Generally, the uncoupling of an output rhythm from the oscillator can be detected in the inconsistency of the phase relationship between them. When the output rhythm is completely coupled with the cell oscillator, it is expected that every cell shows a similar phase relationship.

Figure 5a shows a plot of the peak time difference between *AtCCA1::LUC*+ and *CaMV35S::PtRLUC* against the peak of the *AtCCA1::LUC*+ rhythm for six cells with both rhythms in the same frond (Exp3-1 in Table S1). A positive value of the peak time difference indicates that the phase of *AtCCA1::LUC*+ is advanced compared to that of *CaMV35S::PtRLUC*, e.g., Figure 4b,c. A negative value indicates that the phase of *AtCCA1::LUC*+ is delayed compared to that of *CaMV35S::PtRLUC*, e.g., Figure 4d,e. When the phase relationship is consistent among cells, the peak time difference is plotted horizontally. Instead, the phase relationship between *AtCCA1::LUC*+ and *CaMV35S::PtRLUC* varied greatly among cells, as the time difference was distributed from −12 to 6 h. The inconsistency of the phase relationship was also observed among nine cells, with both rhythms in another frond (Figure S11). Thus, the uncoupling of *CaMV35S::PtRLUC* from *AtCCA1::LUC*+ was clearly shown in cells with both rhythms. This suggests that the *CaMV35S::PtRLUC* rhythm in a cell may not be generated by the cellular circadian oscillator. Although the peak time difference was distributed from −12 to 6 h, the points appeared to be orderly scattered on lines with a slope of −1 on the plot (Figure 5a). This suggests that while the phase of *AtCCA1::LUC*+ varies from cell to cell, the *CaMV35S::PtRLUC* rhythm is in phase among the six cells. Figure 5b shows the peak time distribution of *AtCCA1::LUC*+ rhythm daily (left panel), and its average peak times with the degree of synchronicity (right panel). The average peak time of *AtCCA1::LUC*+ was similar each day, but the degree of synchronicity was lower than 0.55. In contrast, *CaMV35S::PtRLUC* rhythms in the six cells showed similar peak times each day, and the degree of synchronicity was higher than 0.93 (Figure 5c). The higher degree of synchronicity among *CaMV35S::PtRLUC* rhythms than among *AtCCA1::LUC*+ rhythms was also observed in another frond (Figure S11). Thus, it is shown that the *CaMV35S::PtRLUC* rhythms in individual cells are highly synchronous in a frond with asynchronous cellular oscillators.

**Figure 5.**
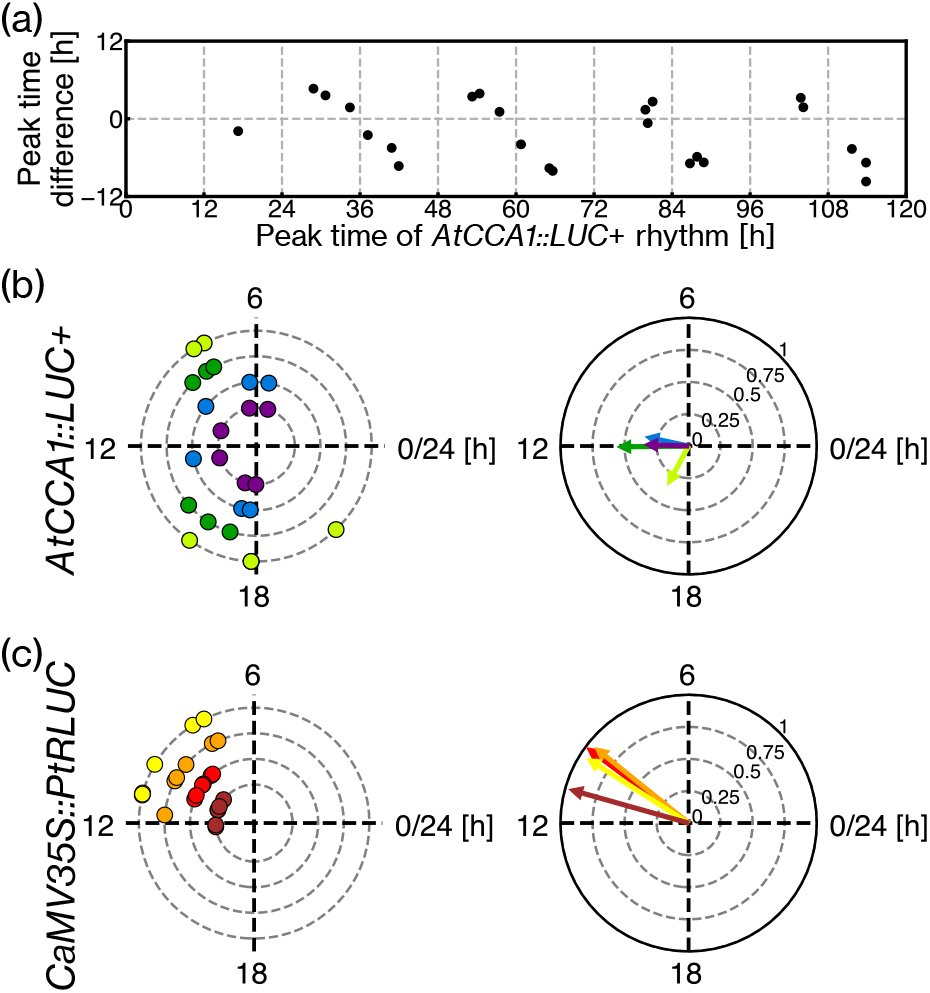
Characterization of phase relationship of cellular bioluminescence rhythms of *AtCCA1::LUC*+ and *CaMV35S::PtRLUC* in the co-transfection experiment. (a) Peak time differences between *CaMV35S::PtRLUC*- and *AtCCA1::LUC*+ rhythms in Exp3-1. Peak time differences between the two reporters for cells showing circadian rhythms (RAE < 0.2) in both reporters (total six cells in this experiment) are plotted. Peak time differences are calculated by the subtraction of a peak time of *AtCCA1::LUC*+ rhythm from that of *CaMV35S::PtRLUC* rhythm, and those ranging from −12 to 12 h are plotted against each peak time of *AtCCA1::LUC*+ rhythm. (b, c) Plots for peak times and their averages/synchronicity of cellular bioluminescence rhythms in Exp3-1. Peak times of the six cells are plotted for *AtCCA1::LUC*+ (b) and *CaMV35S::PtRLUC* (c) on the left. Data display and plotting are the same as in (Figure 4c,e,g), except dots are color-coded by the day of measurement. By regarding individual plots as vectors with a magnitude of 1, the vector sum of the plots for each day is indicated by the arrow of the same color-code on the right. The angle and the magnitude of the vector sum represent the average peak time and the degree of synchronicity, respectively.

In addition to the difference in synchronicity, a difference in free-running periods was observed between *AtCCA1::LUC*+ and *CaMV35S::PtRLUC* rhythms(Figure S12). Out of the 21 cells in the six co-transfection experiments, 17 cells showed a shorter period length in *CaMV35S::PtRLUC* rhythm than in *AtCCA1::LUC*+ rhythm. Furthermore, the correlation coefficient between the two periods was *r* = −0.11, suggesting that the period of *CaMV35S::PtRLUC* rhythm may not be affected by that of the cellular oscillator in individual cells.

In summary, by using the dual-color bioluminescence monitoring system, we observed inconsistencies in the phase relationship, synchronicity, and periodicity between *AtCCA1::LUC*+ and *CaMV35S::PtRLUC* rhythms in the same cells in a frond. Thus, we clearly demonstrate the uncoupling of an output rhythm from the cellular oscillator at the cell level.

## Discussion

The plant circadian clock is a cell-autonomous oscillation system based on the transcription–translation feedback loops, and its time information is output via transcription control (Nagel and Kay, 2012). In a day/night cycle, each cellular oscillator is synchronized to the cyclic environment. Meanwhile, in a constant environment without any cue of day/night signals, those cellular oscillators tend to become desynchronized at the tissue/organ level in the same plant even though the circadian oscillation is maintained in individual cells (Fukuda *et al*., 2007; Fukuda *et al*., 2012; Wenden *et al*., 2012; Muranaka and Oyama, 2016). The heterogeneity of cellular oscillators in the same tissue is an important force for desynchronization (Muranaka and Oyama, 2016; Okada *et al*., 2017; Muranaka and Oyama, 2018). It would be impossible to analyze the circadian system of individual cells in an asynchronous state by observing whole tissues/organs because the output should be arrhythmic at the tissue/organ level. Simultaneous measurement of cellular behaviors of the oscillator and output in the same cells could be a powerful tool to solve this problem. We developed a dual-color bioluminescence monitoring system that can monitor the luminescence of two reporters simultaneously at a single-cell level, and we clearly demonstrated the uncoupling between the cellular oscillator and output phenomenon in a duckweed plant in an asynchronous state. Here, we introduced *AtCCA1::LUC*+, which reports the behavior of cellular oscillator with yellow-green luminescence, and *CaMV35S::PtRLUC*, which shows circadian output with red luminescence (Muranaka *et al*., 2015; Nishiguchi *et al*., 2015; Muranaka and Oyama, 2016). By quantitative analysis of the luminescence behaviors of these reporters in co-transfected cells, we obtained the following two pieces of evidence of the uncoupling of the *AtCCA1::LUC*+ and *CaMV35S::PtRLUC* rhythms: (1) irrespective of the stability (presence or absence) of the *CaMV35S::PtRLUC* rhythm, *AtCCA1::LUC*+ showed a robust rhythm basically in each cell; and (2) cellular *CaMV35S::PtRLUC* rhythms, which were unstable and noisy, showed a relatively high synchrony even in the frond with a low synchrony of cellular *AtCCA1::LUC*+ rhythms. The uncoupling between these two rhythms could be supposed on the bases of the different proportions of rhythmic cells in the single-reporter transfection experiments (Figure 4a). However, it would be difficult to quantitatively analyze the relationship between these two rhythms by using data from the single-reporter transfection experiments because of the uncontrollable synchronicity of cellular circadian oscillators in plants without experiencing day/night cycles (Wenden *et al*., 2012; Muranaka and Oyama, 2016).

In this study, we focused on the cellular behaviors of *AtCCA1::LUC*+ and *CaMV35S::PtRLUC* luminescence in a frond in an asynchronous state to exhibit advantages of the dual-color bioluminescence monitoring system. As shown in Figure S2, the circadian rhythmicity of *CaMV35S::PtRLUC* luminescence seemed dependent on the state of synchronization in the frond. The degree of synchrony of cellular circadian rhythms in plants/organs is an important factor for the entire circadian system (Masuda *et al*., 2017; McClung, 2019). In general, a high degree of synchrony is required for the physiological function of timekeeping in plants. Interestingly, the entrainment manners of cellular oscillators in *Arabidopsis* are presumed to be dependent on the degree of synchrony; the circadian rhythm of plants in an asynchronous state is more sensitive to a dark pulse for entrainment than that of plants in a synchronous state (Masuda *et al*., 2017). The circadian behavior of *CaMV35S::PtRLUC* luminescence seems to be a synchrony-dependent phenomenon and may be related to other synchrony-dependent phenomena. The mechanism that gives *CaMV35S::PtRLUC* luminescence circadian rhythmicity is unknown. The uncoupling of the rhythm from the cellular oscillator suggests that *CaMV35S::PtRLUC* may not be directly controlled by the transcriptional and translational feedback loops of clock-related genes, and PtRLUC may be expressed without circadian rhythmicity. In fact, the *CaMV35S* promoter is known as a constitutive promoter and its expression did not show a circadian rhythm in tobacco (Millar *et al*., 1992). Similarly to *CaMV35S::PtRLUC, ZmUBQ::LUC*+ showed a circadian luminescence rhythm in various duckweed plants (Muranaka *et al*., 2015). *ZmUBQ* is known as a constitutive promoter for monocotyledonous plants (Christensen *et al*., 1992; Miwa *et al*., 2006). The circadian rhythms of these reporters observed in duckweed plants might be generated at the level of post-transcriptional regulation. It is known that many gene products of plants are circadianly regulated at various post-transcriptional steps (McClung, 2019). The intracellular concentrations of substrates of luciferase are possible factors that can be responsible for the circadian rhythms of these bioluminescent reporters. Firefly luciferase catalyzes the ATP-dependent oxidation of luciferin (Van Leeuwen *et al*., 2000). The circadian regulation of photosynthesis has been analyzed in many plant species (Dodd *et al*., 2014). The production of ATP and oxygen is dependent on photosynthetic activity and changes in the intracellular concentrations of these substrates of luciferase may be in a circadian manner. It was reported that the ATP accumulation of *Lemna gibba* showed a circadian rhythm under constant light after light/dark cycles (Kondo and Nakashima, 1979). These molecules can diffuse through the apoplast or symplast, and their intracellular concentrations may be spatially equalized in a region of the plant. Such diffusive molecules could be a mediator of cellular circadian rhythms that are uncoupled from the cellular oscillator but are locally synchronized with each other. In addition to these molecules derived from photosynthesis, it has been shown that there is a circadian rhythm in the absorption of ions, such as potassium and magnesium, and these diffusive ions may be involved in the circadian rhythmicity of the luciferase reaction or luciferin uptake (Kondo and Tsudzuki, 1978; Kondo, 1983; Feeney *et al*., 2016). The circadian rhythms of these low-molecular and diffusive substances appear to be coupled with the timekeeping of various metabolic pathways in plants; the temporal coordination of these pathways is important. Luciferase reporters under the control of a constitutive promoter may be a useful tool to spatiotemporally analyze the timekeeping system.

In this study, we used the firefly LUC+ as a short-wavelength luciferase in the dual-color bioluminescence monitoring system. Almost all bioluminescent transgenic plants carry this luciferase and its applications for plant science are widespread (Van Leeuwen *et al*., 2000). They are preferably used in various analyses of gene expression dynamics. The enzymatic reaction of the firefly luciferase depends on the reaction conditions and its emission spectrum is altered by pH changes. Based on our single-cell transfection experiments using the firefly luciferase, it seems that the higher the expression level, the longer the emission wavelength (Figure S4). The enzyme reaction conditions may change due to the excessive accumulation of luciferase. In contrast, PtRLUC, which we used as a long-wavelength luciferase, appeared to present an emission spectrum independently of its expression level (Figure S3b). In the dual-color bioluminescence monitoring system, the selection of filters to separate luminescence between the two luciferases is critical. Ideally, the green- and red-pass filters should transmit luminescence only from LUC+ and PtRLUC, respectively, and the band-pass range of each filter should include the wavelength of the maximum emission of each luciferase. Because of the overlap of the emission spectra between LUC+ and PtRLUC (Figure 1b), we had to use filters deviated from the ideal and had to exclude many cells with poor proportions of luminescence intensities between two reporters for the following analysis. A combination of luciferases with a smaller overlap of their emission spectra is preferable. Using a luciferase with maximum emission at a shorter wavelength than firefly LUC+ (λ_max_ = ~560 nm) is an option; there are several available insect-derived luciferases (and their derivatives) in this category (λ_max_ < ~550 nm) (Viviani, 2002). Blue-emitting luciferases (λ_max_ < 490 nm) such as *Renilla* luciferase and bacterial luciferases, would be a spectrally good option; however, their luminescence reactions are completely different from those of insect luciferases (Shimomura, 2006). Treating plants with the same substrate, luciferin, for insect-derived luciferases is critical to ensure the homogeneous enzyme reaction conditions for the two luciferases in cells. In dual-color bioluminescence monitoring experiments previously reported for mammals and cyanobacteria, insect-derived luciferases in the shorter wavelength category were utilized (Kitayama *et al*., 2004; Nakajima *et al*., 2004; Nakajima *et al*., 2005; Ogura *et al*., 2005; Kwon *et al*., 2010; Nishide *et al*., 2018).

The dual-color bioluminescence monitoring system at a single-cell level is useful to deeply analyze the relationship (e.g., correlation) between two gene expression behaviors. Especially, stochastic phenomena found in gene expression (reporter activities) are interesting targets for analysis using this monitoring system (Kærn *et al*., 2005). In this study, the circadian rhythmicity of *CaMV35S::PtRLUC* and phases of *AtCCA1::LUC*+ rhythms are such stochastic phenomena. It has been difficult to approach the underlying regulatory networks of various stochastic phenomena accompanying large cell-to-cell variation in plants. The dual-color bioluminescence monitoring system at a single-cell level could become an experimental tool to directly analyze the behavior of components in the networks. While we analyzed the luminescence behaviors of two reporters at the single-cell level, similar analyses on stochastic phenomena found in larger spatial regions, such as those within and between individuals, can be performed using transgenic plants expressing two bioluminescent reporters. It is known that under constant conditions, transgenic *Arabidopsis* plants carrying the *AtCCA1::LUC* reporter exhibit a variety of spatiotemporal patterns of luminescent rhythms with significantly different phases even within the same organ (Fukuda *et al*., 2007; Fukuda *et al*., 2012; Wenden *et al*., 2012). By combining another bioluminescent reporter for a clock or output gene, the free-running circadian system at the tissue, organ, and plant body levels should be deeply analyzed for its regulatory networks. In addition, by combining a tissue-specific or age-dependent reporter with a circadian reporter, the dynamics of the circadian system in growing plants can be effectively analyzed. Furthermore, the dual-color bioluminescence monitoring system at a single-cell level can be used together with genetic manipulation at a single-cell level. It was reported that the co-transfection of effectors for the overexpression, knock-down (by RNAi), and knock-out (by CRISPR/Cas9) of clock genes with a circadian bioluminescent reporter gave rise to alterations in the circadian behavior of transfected cells in duckweed and *Arabidopsis* (Miwa *et al*., 2006; Serikawa *et al*., 2008; Okada *et al*., 2017; Kanesaka *et al*., 2019). By analyzing the effects of genetic manipulation by effectors on the luminescence behavior of two circadian reporters in individual cells, the structure of the circadian system can be elucidated.

In this study, we used *L. minor* as the plant material. Since duckweeds are generally small and flat, they have been used for physiological experiments in the laboratory (Stomp, 2005; Muranaka and Oyama, 2018). In studies on plant circadian rhythms, physiological rhythms, such as CO_2_ output and ion uptake, were well characterized by using duckweeds (Miyata and Yamamoto, 1969; Hillman, 1970; Kondo and Tsudzuki, 1978). Furthermore, functional analysis of the clock-related genes in *Lemna* plants revealed the conservation of their gene functions in plants (Miwa *et al*., 2006; Serikawa *et al*., 2008; Okada *et al*., 2017). Thus, knowledge on circadian systems, including clock-related genes, has been accumulated for duckweed. Duckweed plants show a large diversity of structural complexity among species. The number of shoot apical meristems, the existence of roots, and the development of tracheid tissue are dependent on the species/genus (Les *et al*., 1997). In *Arabidopsis*, the shoot apex and vascular tissues are regarded somewhat as pacemakers in the plant body (Fukuda *et al*., 2007; Fukuda *et al*., 2012; Endo *et al*., 2014; Takahashi *et al*., 2015). Their roles in the circadian system of plants will be approached by analyzing various duckweed plants. Basic circadian properties of various duckweed species have been already analyzed by a bioluminescence monitoring system with particle bombardment (Muranaka *et al*., 2015; Isoda and Oyama, 2018). The bioluminescence monitoring system at a single-cell level is also available in *Arabidopsis* leaves, and the circadian rhythms of individual cells can be detected (Kanesaka *et al*., 2019). In future, the dualcolor bioluminescence monitoring system will be applied for the study of the circadian system in *Arabidopsis*.

Recently, methods for the stable transformation of *L. minor* have been established (Chhabra *et al*., 2011). Thus, spatiotemporal analysis for circadian rhythm can be performed using a transgenic duckweed carrying a circadian bioluminescent reporter. Because duckweed grows flat on a liquid surface, it is easier to perform the long-term high-resolution monitoring of proliferating plants (Muranaka and Oyama, 2018). By transiently introducing a circadian bioluminescent reporter by particle bombardment into the transgenic duckweed, it will be possible to simultaneously observe the general behavior of circadian rhythm in the fronds and circadian behavior in individual cells. Analyzing combinations of various circadian reporters will elucidate complex circadian behavior in plants.

## Experimental procedures

### Plant material and growth conditions

*Lemna minor* 5512 was maintained in NF medium with 1% sucrose under constant light conditions in a temperature-controlled room (25 ± 1 °C) as previously described (Muranaka *et al*., 2015). The white light (~20 μE m^−2^ s^−1^) was supplied by fluorescent lamps (FLR40SEX-W/M/36-HG, NEC, Tokyo, Japan). Plants were grown on 60 ml of the medium in 200-ml Erlenmeyer flasks plugged with cotton. New stock cultures were made weekly, and well-grown plants were used for experiments. Treatment of *Lemna minor* plants with light/dark cycles was carried out in an incubator (MIR-153, Panasonic, Tokyo, Japan).

### Luciferase reporter transfection by particle bombardment

As circadian bioluminescent reporters, we used *pUC-AtCCA1::intron-LUC*+ (*AtCCA1::LUC*+), *pUC18-CaMV35S::intron-luc*+ (*CaMV35S::LUC*+) (Muranaka *et al*., 2015), and *pUC18-CaMV35S::PtRLUC (CaMV35S::PtRLUC). pUC-AtCCA1::intron-LUC*+ is a modified version of *pUC-AtCCA1::LUC*+, which has been described previously (Muranaka *et al*., 2013), with an intron sequence inserted at 148 bp from the start codon of the luciferase gene. The *intron-LUC*+ was described previously by Muranaka *et al*., (2015). *pUC18-CaMV35S::PtRLUC* is a reporter driven by the *CaMV35S* promoter which expresses PtRLUC, a red-shift mutant (S246H/H347A) of Emerald luciferase (ELuc) from Brazilian click beetle (*Pyrearinus termitilluminans*) (Nishiguchi *et al*., 2015). Reporter constructs were introduced into *L. minor* plants by particle bombardment as described previously (Muranaka *et al*., 2015). In both single- and dual-reporter experiments, the amounts of plasmid DNA for *AtCCA1::LUC+, CaMV35S::PtRLUC*, and *CaMV35S::LUC*+ were 2 μg, 0.2 μg, and 0.2 μg, respectively.

### Fluorescence imaging

A fluorescent reporter (*CaMV35S::GFP-h*; Nakano *et al*., 2009) was transfected to duckweed plants and GFP fluorescence-positive cells were observed under confocal microscope (LSM510-META; Carl Zeiss, https://www.zeiss.com) as described previously (Muranaka *et al*., 2013; Isoda and Oyama, 2018).

### Bioluminescence monitoring

Single-cell bioluminescence imaging was basically carried out as described previously (Muranaka and Oyama, 2016; Muranaka and Oyama, 2020), with modifications for dual-color bioluminescence and multi-sample monitoring. Two colonies of *L. minor* transfected with a reporter(s) were transferred to a 35-mm polystyrene dish with 3 ml of growth medium containing 0.5 mM D-luciferin. Two sample dishes (i.e., four colonies) were monitored in an experiment. Sample dishes were set on a handmade motor-driven stage with rotation and z-axis stages (ARS-6036-GM and ALV-902-HP; Chuo Precision Industrial Co., Tokyo, Japan) in a lightproof box in an incubator (25 ± 1°C) (Muranaka and Oyama, 2020). The stage rotated to change the position of the samples underneath the lens of an EM-CCD camera (ImagEM C9100-13; Hamamatsu Photonics, Hamamatsu, Japan) or an end-emitting fiber optics guiding LED light (approximately 30 μE m^−2^ s^−1^; RFB2-20SW; CCS Inc, Kyoto, Japan). A handmade motor-driven filter changer with a rotation motor (ARS-4036-GM; Chuo Precision Industrial Co.) was set between the sample and the lens. A green-pass filter (PB0530/080; ø49.7 mm; Asahi Spectra, Tokyo, Japan) and red-pass filter (PB0630/040; ø49.7 mm; Asahi Spectra) were set on slots in the filter changer, and a slot was left empty to capture unfiltered images. By controlling this imaging system using PC software (HOKAWO; Hamamatsu Photonics), a series of bioluminescence images for a sample dish were automatically captured every 1 h. Before image capturing, we allowed at least a 4 min dark period for autofluorescence decay. After the dark period, images were captured for each filter condition: unfiltered, green-pass filtered, and red-pass filtered, with exposure times of 20 s, 120 s, and 30 s, respectively. The focus for each filter condition was optimized by adjusting the distance between the sample and the lens by driving the z-axis motor of the stage. During the monitoring, the focal adjustment was performed as follows: from the reference position (unfiltered bioluminescence image in focus), 1.0-mm downward for the green-pass filter and 1.8-mm downward for the red-pass filter. To remove cosmic ray spikes, two sequential images at a time for each filter condition were captured and the minimum value for each pixel was used for analysis. These exposure procedures accompanied an approximately 18-min dark period per hour.

Bioluminescence monitoring at a whole plant level (Figure S2) was performed as previously described by Muranaka *et al*. (2015).

### Luminescence spectra

The luminescence emission of *L. minor* transfected with *CaMV35S::LUC*+ or *CaMV35S::PtRLUC* by particle bombardment was filtered by a series of optical band-pass filters with a narrow band width (half width 10 nm; Asahi Spectra), and the intensity of the filtered luminescence was obtained from the image of a luminous frond captured by the EM-CCD camera. Intensities normalized to the maximum were plotted.

### Time series analysis

The bioluminescence intensity of each luminescent spot was measured as the integrated density of the region of interest (size, 6 × 6 pixels). Image analysis and postprocessing were carried out with ImageJ (http://rsbweb.nih.gov/ij/) as previously described by Muranaka and Oyama (2016) and Muranaka and Oyama (2020). The following analysis was implemented by Python 3.7.4 (NumPy 1.16.5, SciPy 1.3.1). The terms and symbols for luminescence intensities are defined as follows.

Unfiltered luminescence intensity (*L*), measured value of luminescence captured without a filter; Green luminescence intensity (*L_g_*), measured value of luminescence transmitted through the green-pass filter; Red luminescence intensity (*L_r_*), measured value of luminescence transmitted through the red-pass filter; LUC+ luminescence intensity (*L_LUC+_*), LUC+ luminescence value calculated from *L_g_* and *L_r_*; PtRLUC luminescence intensity (*L_PtRLUC_*), PtRLUC luminescence value calculated from *L_g_* and *L_r_*. The luminescence intensity of each luminous spot was quantified by the following equation according to the manufacturer’s instructions (Muranaka and Oyama, 2020): Number of photons = (signal intensity – background intensity) × 5.8/(1200 × 0.9). As explained in Figure S3, *L_LUC+_* and *L_PtRLUC_* were calculated from *L_g_* and *L_r_* using the following equation:

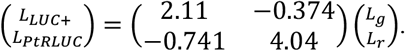

The RAEs of bioluminescence traces were estimated using the FFT-NLLS method as previously described (Muranaka and Oyama, 2016), using data in the time range of 24–96 h. The peak times of bioluminescence traces were estimated by a local quadratic curve fitting (Muranaka and Oyama, 2016). Time series data were smoothed with the 12-h moving average, and peak positions were roughly identified as local maxima. Precise positions of peaks were estimated by a local quadratic curve fitting using the original time series data. The width of the local fitting area was set to 9 h.

## Acknowledgments

We thank Dr. Masaaki Morikawa for providing us with *Lemna minor* 5512. This work was supported in part by the Japan Society for the Promotion of Science KAKENHI (Grant numbers 25650098, 17KT0022, and 19H03245), the Japan Science and Technology Agency (JST) ALCA to TO, and KAKENHI (Grant number 20K06342) to SI, and KAKENHI (Grant number 24-1530) to TM.

## Author contributions

EW, TM, SI, and TO conceived and designed the research and experiments. EW, TM, SI, and TO constructed the dual-color monitoring system. EW performed bioluminescent reporter experiments and data analyses. MI performed fluorescent reporter experiments. EW and TO wrote the manuscript.

## Supporting Information for

### Legends for supporting information

**Figure S1.**
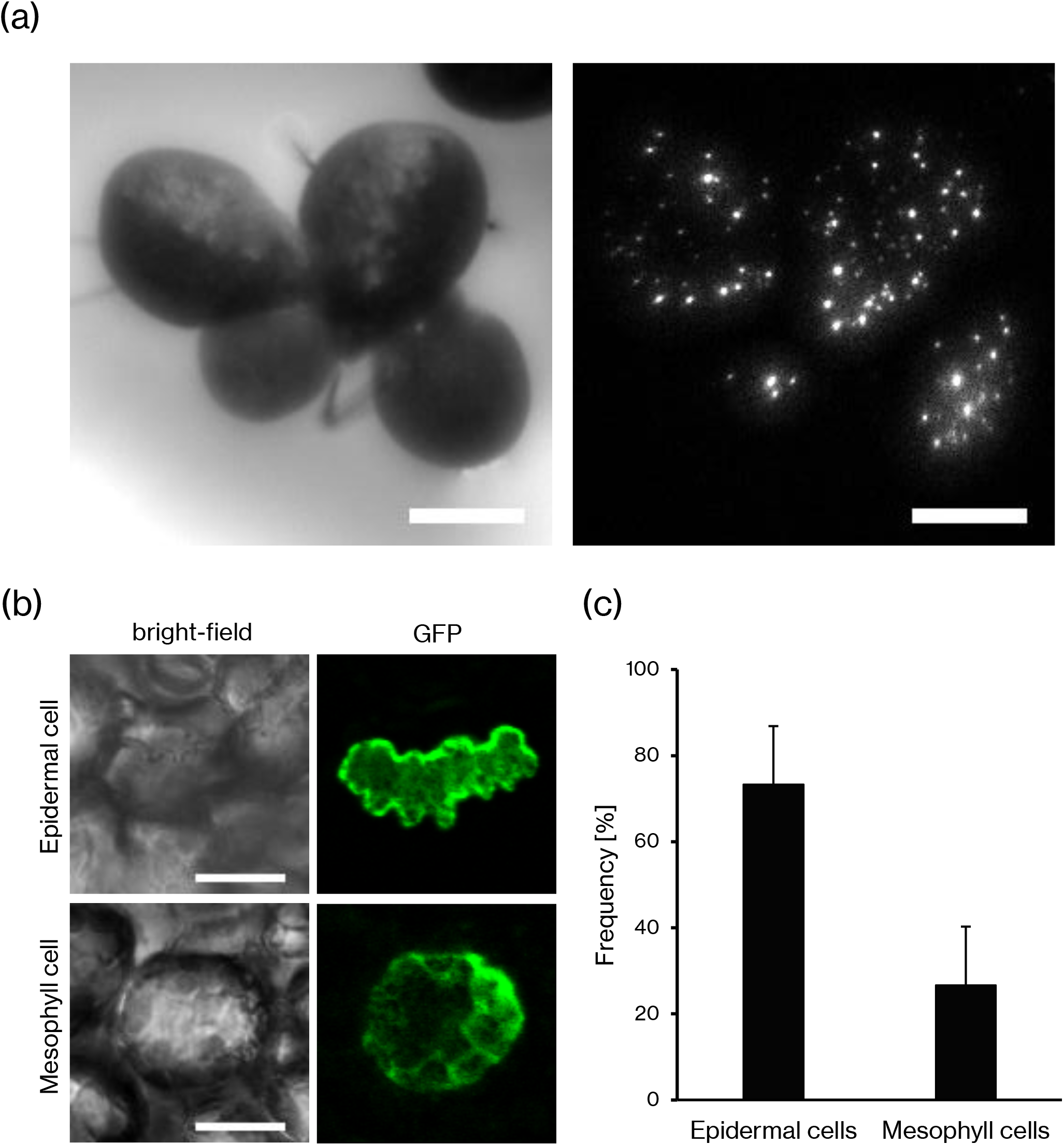
Transfection experiments of *Lemna minor* by particle bombardment. (a) A bright-field image of *L. minor* transfected with the bioluminescence reporter *CaMV35S::PtRLUC* (left), and a luminescence image captured 24 h after particle bombardment (right). (b) An epidermal cell (upper) and a mesophyll cell (lower) that were transfected with the fluorescence reporter gene *CaMV35S::GFP-h*. A bright-field image (left) and fluorescence image (right) are shown. (c) The 1252 fluorescent cells in three independent experiments were observed and classified into epidermal or mesophyll cells and their percentages are shown. Error bars represent the SD. Scale bars in (a) and (b) are 2 mm and 20 μm, respectively.

**Figure S2.**
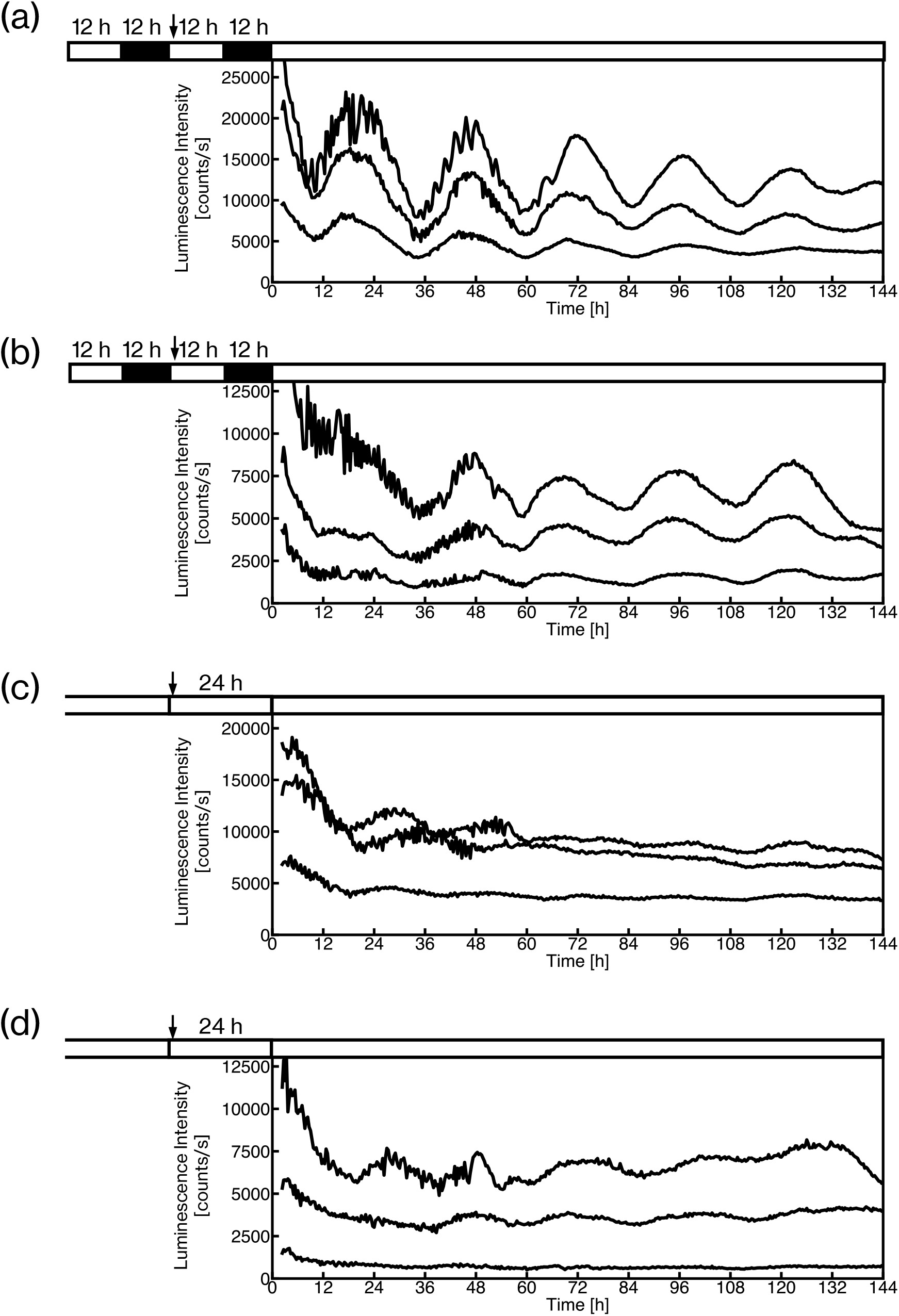
Time series data at a whole plant level for *Lemna minor* transfected with *CaMV35S::LUC*+ or *CaMV35S::PtRLUC*. *CaMV35S::LUC*+ (a, c) or *CaMV35S::PtRLUC* (b, d) was introduced into *L. minor* by particle bombardment; these plants were monitored under constant light conditions by using the luminescence dish-monitoring system with a photomultiplier tube. Three samples were monitored for each condition. The experimental schedule is shown at the upper part of each graph. Black and open bars indicate dark and light conditions, respectively. The timing of gene transfection is shown by an arrow. Duckweed plants had been maintained under light/dark conditions (a, b) or constant light conditions (c, d) before they were used for the experiments.

**Figure S3.**
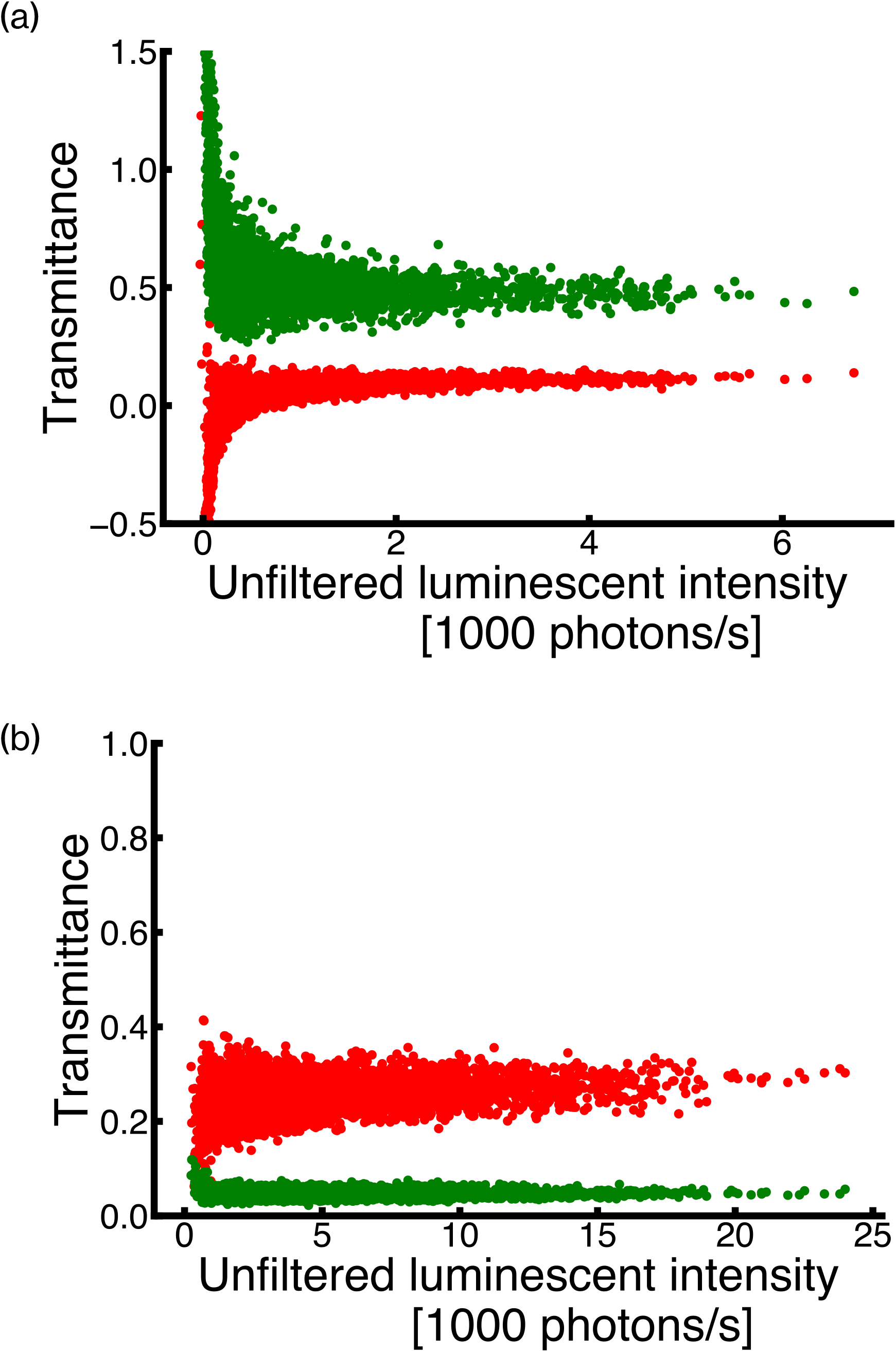
Scatter plots of transmittance versus unfiltered luminescence intensities in single-reporter transfection experiments. Transmittance of LUC+ (a) and PtRLUC (b) for green-pass (green dots) and red-pass filters (red dots) are plotted against unfiltered luminescence intensities. Time series data of the luminescence intensities obtained in Exp1-1 and Exp2-1 were used. Twenty-eight (LUC+) and 38 (PtRLUC) luminous spots with 149 time series data points (149 h) were calculated for transmittance. Data including one or more pixels with a saturated signal in the spot area for one or more filter conditions were excluded. The total data numbers are 4151 (LUC+) and 5558 (PtRLUC).

**Figure S4.**
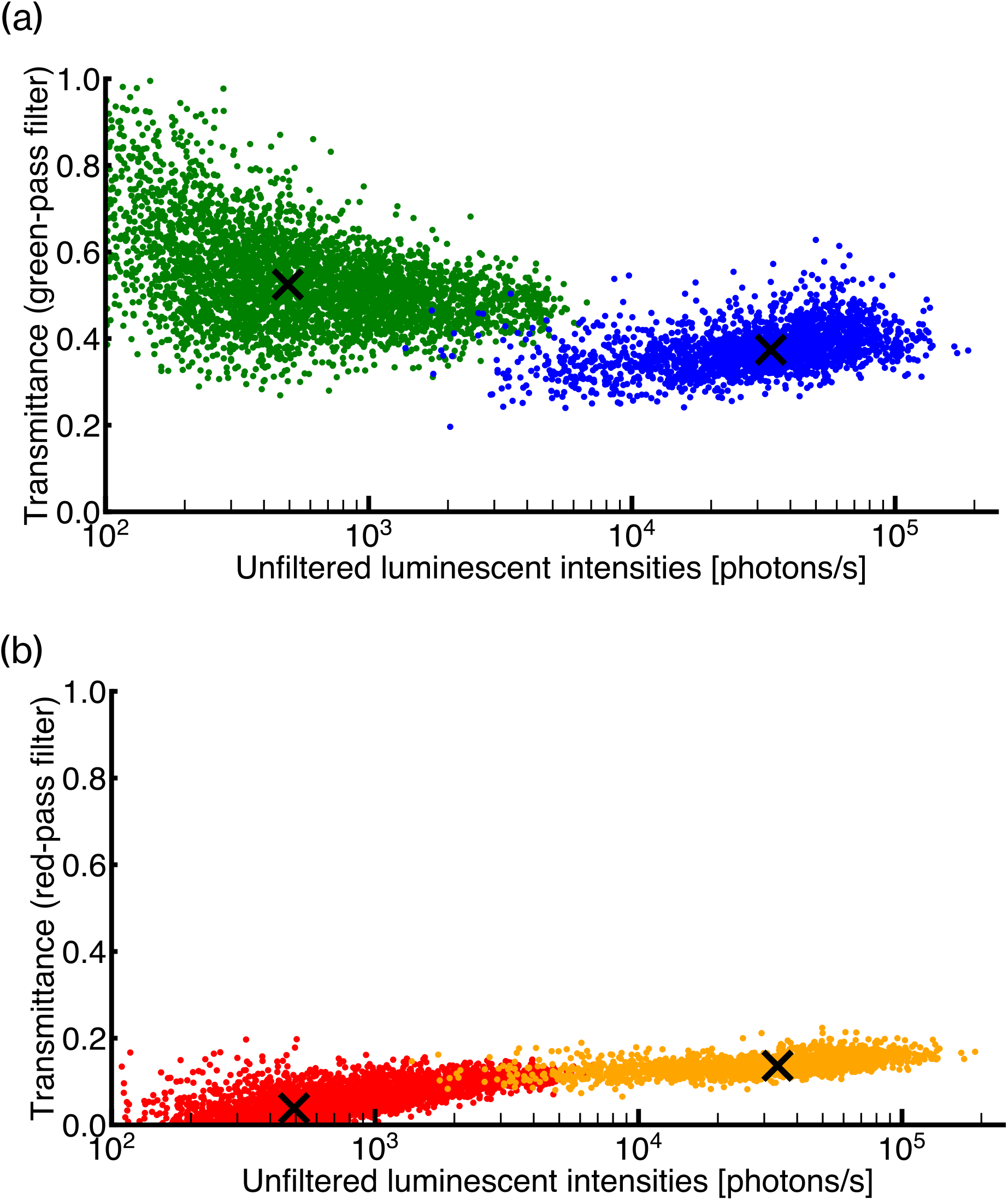
Comparison of transmittance of filters between *AtCCA1::LUC*+ and *CaMV35S::LUC*+. Transmittance for each data point in the time series of luminescence intensities is plotted for *AtCCA1::LUC*+ (Exp1-1, total 4151 spots) and that for luminous spots in snapshot images is plotted for *CaMV35S::LUC*+ (total 2147 spots). (a) Transmittance of *AtCCA1::LUC*+ (green dot) and *CaMV35S::LUC*+ (blue dot) for the green-pass filter is plotted against unfiltered luminescence intensities. (b) Transmittance of *AtCCA1::LUC*+ (red dot) and *CaMV35S::LUC*+ (orange dot) for the red-pass filter are plotted against unfiltered luminescence intensities. The cross mark represents the median for each condition.

**Figure S5.**
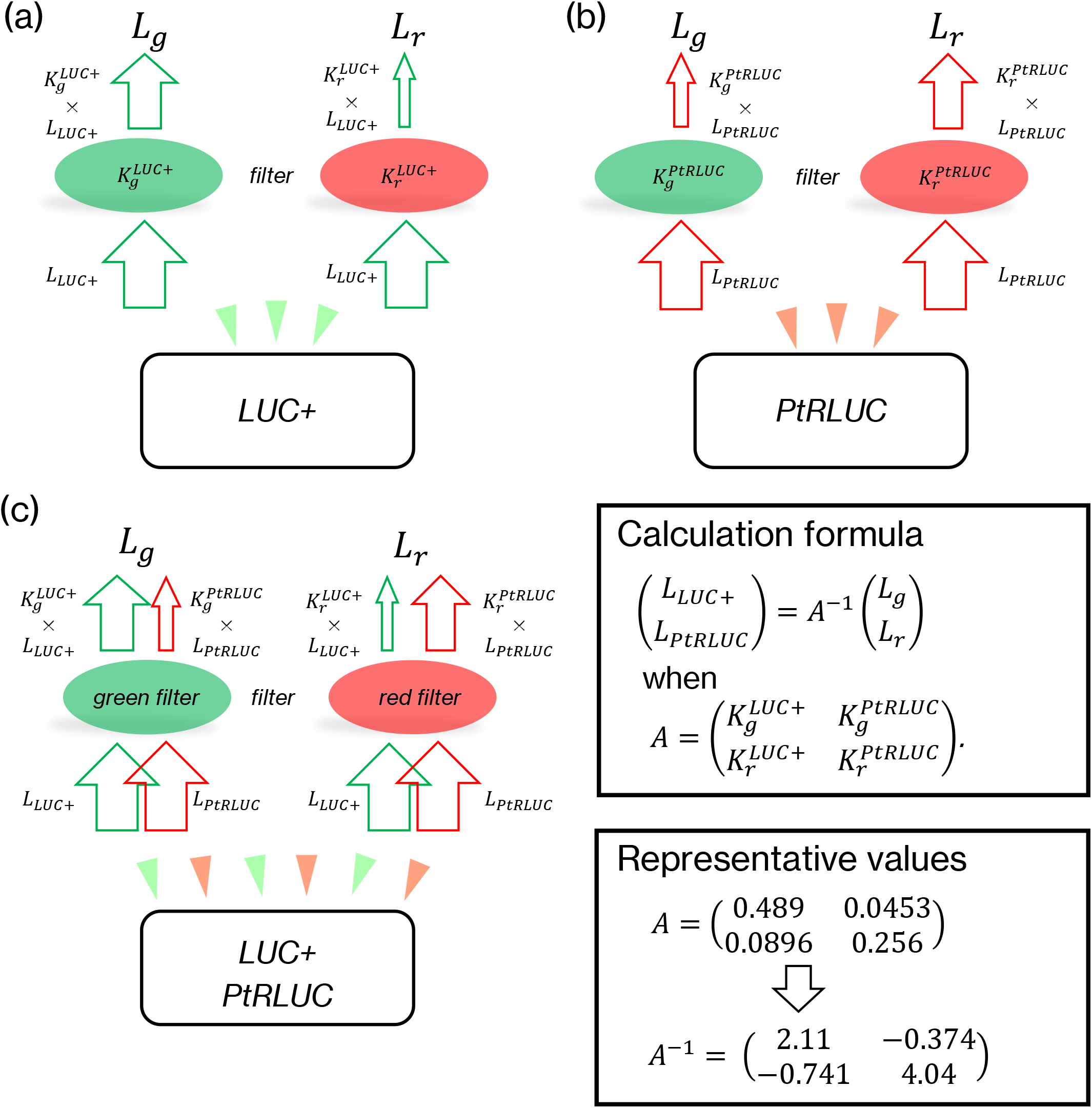
Schematic drawing for the calculation of luminescence intensities of LUC+ and PtRLUC. (a) Transmittance of green-pass filter and red-pass filter for LUC+ luminescence are represented as 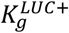 and 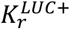, respectively. In the single-reporter transfection experiment, 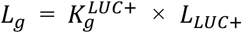, where *L_g_* is luminescence transmitted through the green-pass filter and *L_LUC+_* is LUC+ luminescence, as mentioned in the ‘Experimental procedures’ section. Similarly, 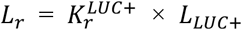. (b) Transmittance of the green-pass filter and red-pass filter for PtRLUC luminescence are represented as 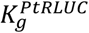 and 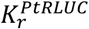, respectively. Similarly to (a), in the single-reporter transfection experiment, 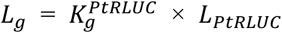 and 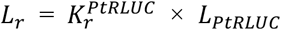. (c) In the co-transfection experiment, 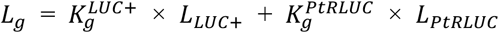 and 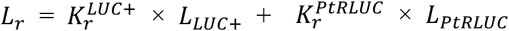. The luminescence intensities from the two luciferases in the co-transfection experiment are calculated with inverse matrix *A^−1^* as in the box for calculation formula. In our experiments, we used the matrix elements described in the box for representative values.

**Figure S6.**
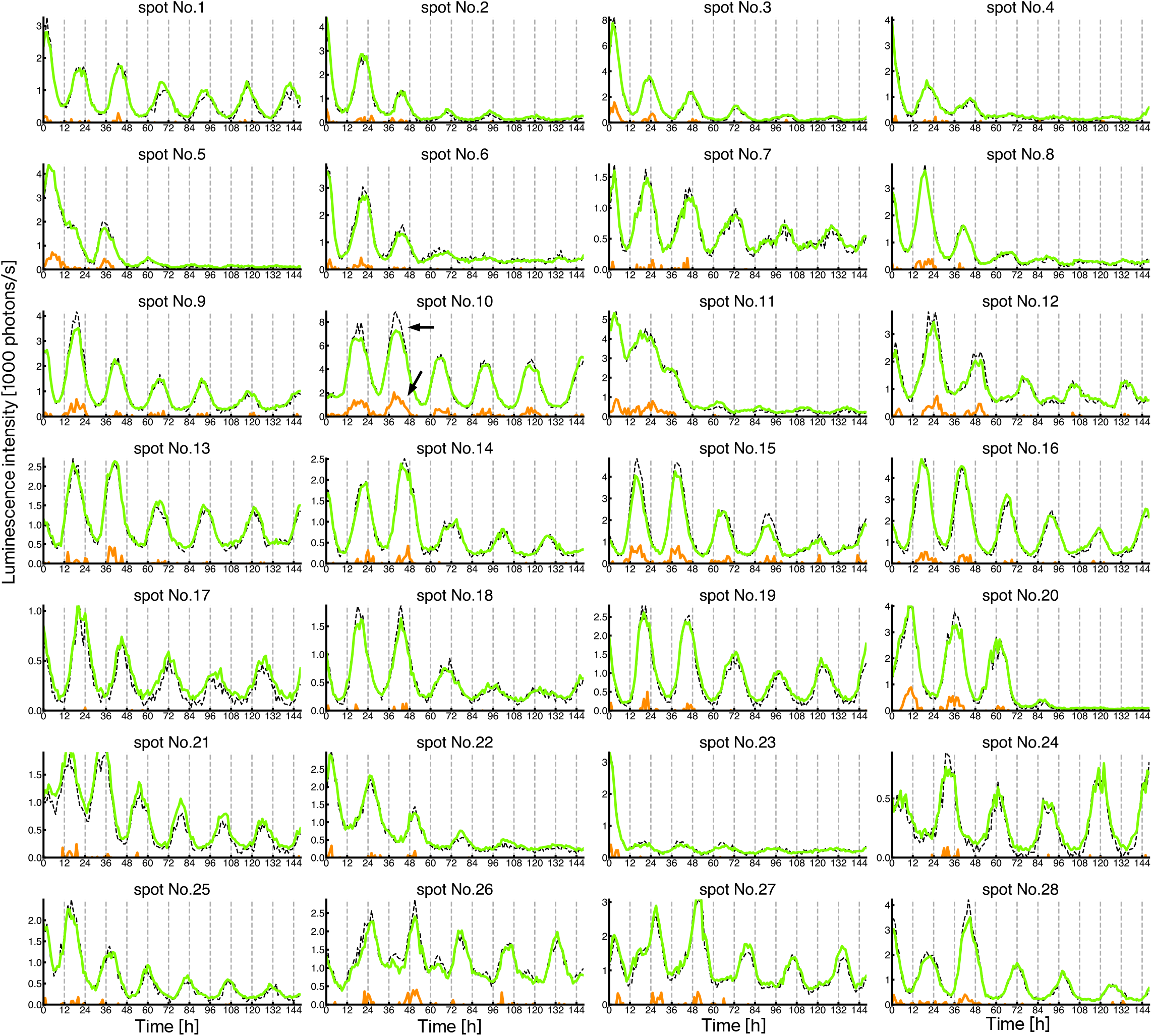
Reconstruction of time series of luminescence intensities of luminous spots for *AtCCA1::LUC*+ in a single-reporter transfection experiment. Luminescence intensities of LUC+ (light-green line) and PtRLUC (orange line) were reconstructed using the time series data of filtered luminescence intensities for each cell in Exp1-1. A dashed line in each graph represents the original time series of unfiltered luminescence intensities. The arrows in spot no. 10 show an example where the *L_LUC+_* and *L_PtRLUC_* values are underestimated and overestimated, respectively.

**Figure S7.**
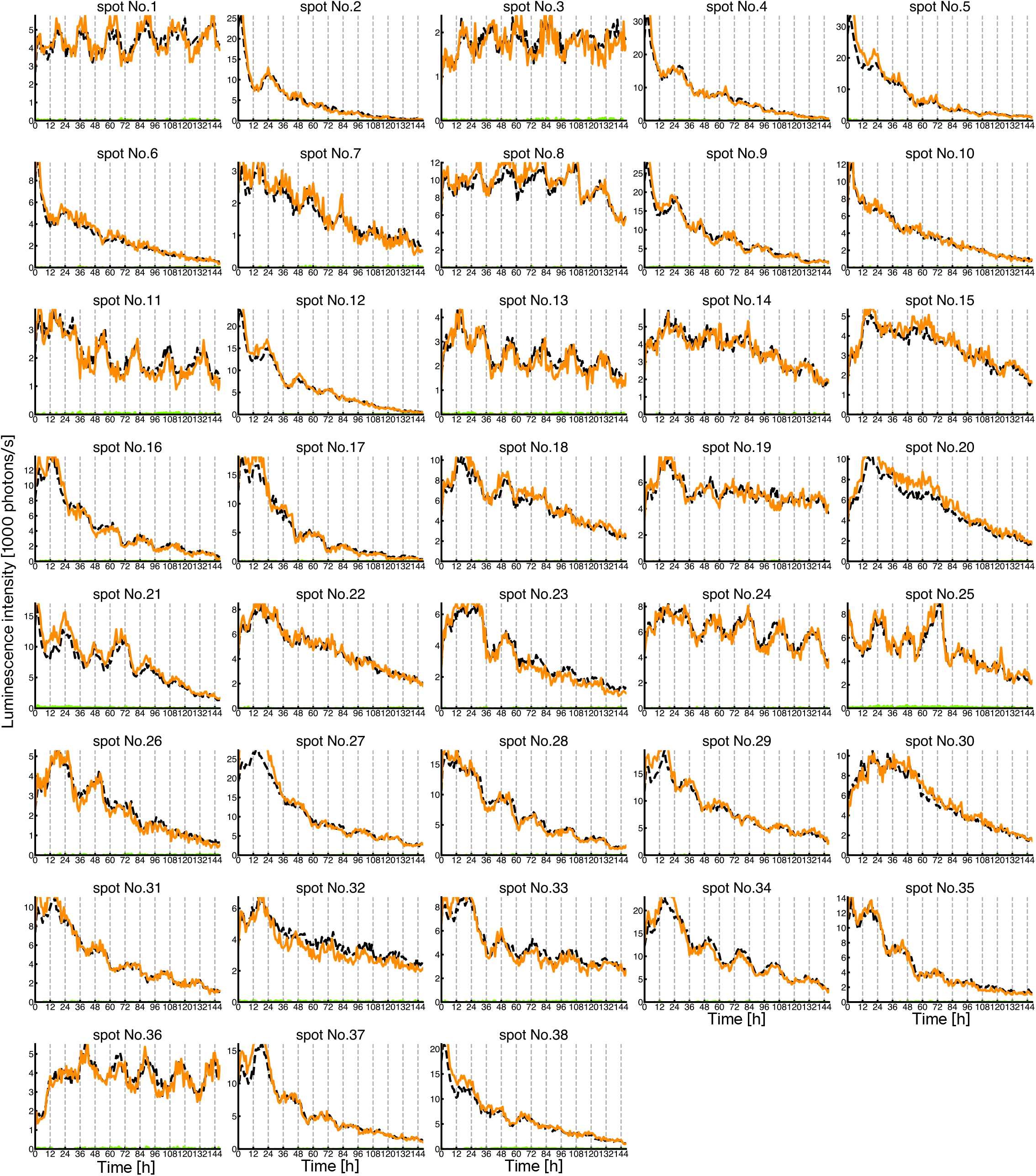
Reconstruction of time series of luminescence intensities of luminous spots for *CaMV35S::PtRLUC* in a single-reporter transfection experiment. Luminescence intensities of PtRLUC (orange line) and LUC+ (light-green line) were reconstructed using the time series data of filtered luminescence intensities for each cell in Exp2-1. A dashed line in each graph represents the original time series of unfiltered luminescence intensities. The time series of spot nos. 2, 4, 5, 9, 17, 27, 28, 29, 34, and 38 include data points with saturated signals in the spot area early in the luminescence measurement. The estimated luminescence of LUC+ is near the baseline and almost invisible in each graph.

**Figure S8.**
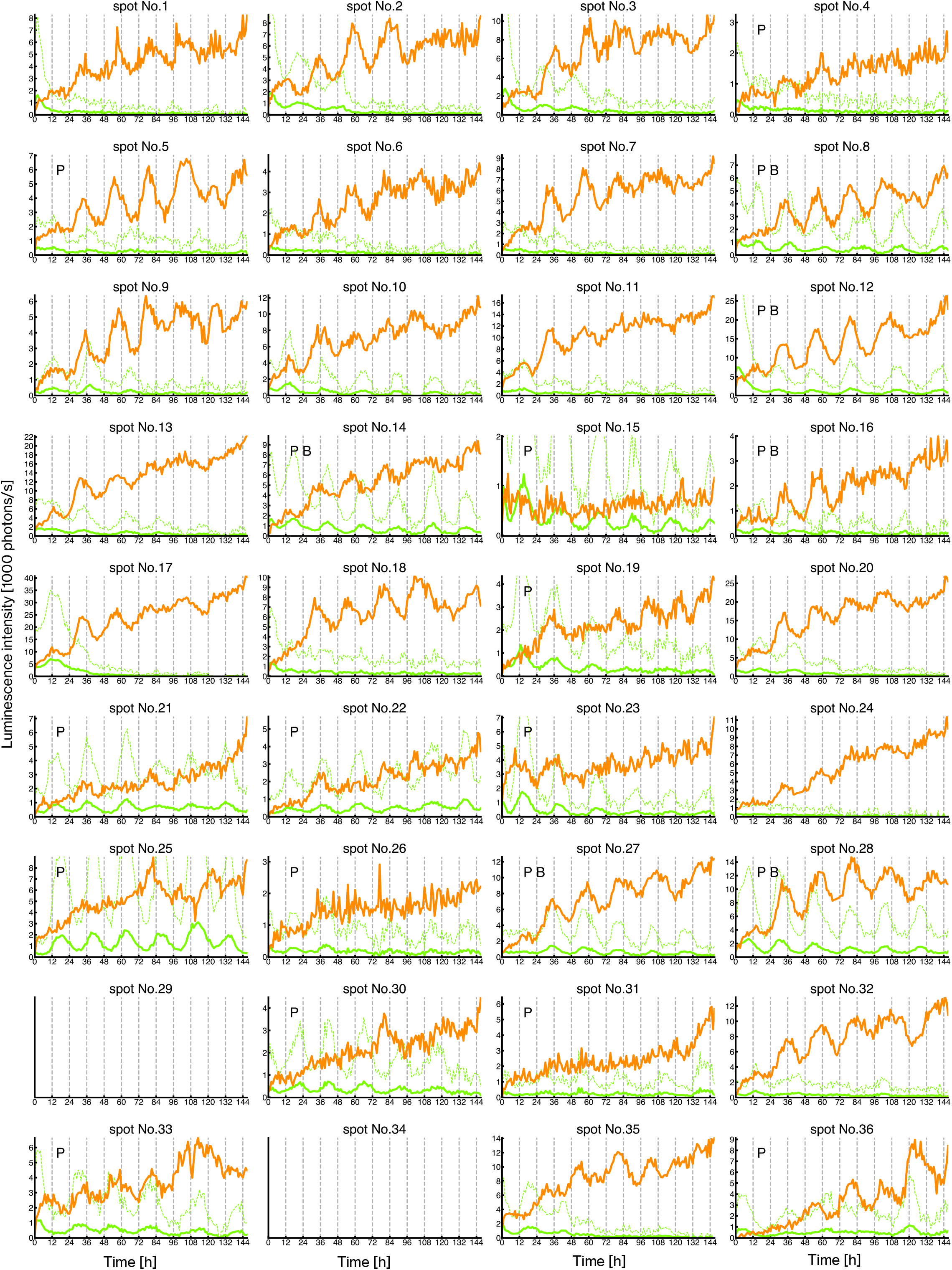
Reconstruction of time series of luminescence intensities of luminous spots in a co-transfection experiment. Luminescence intensities of LUC+ (light-green line) and PtRLUC (orange line) were reconstructed using the time series data of filtered luminescence intensities for each cell in Exp3-1. A dashed light-green line in each graph represents the five-fold magnification of LUC+. P and B in the graph represent analyzable spots and cells showing the circadian rhythms of both reporters, respectively (refer to Figure S9). At many time points, spot areas of spot no. 29 and spot no. 34 included saturation signals, and were excluded (refer to Figure S9).

**Figure S9.**
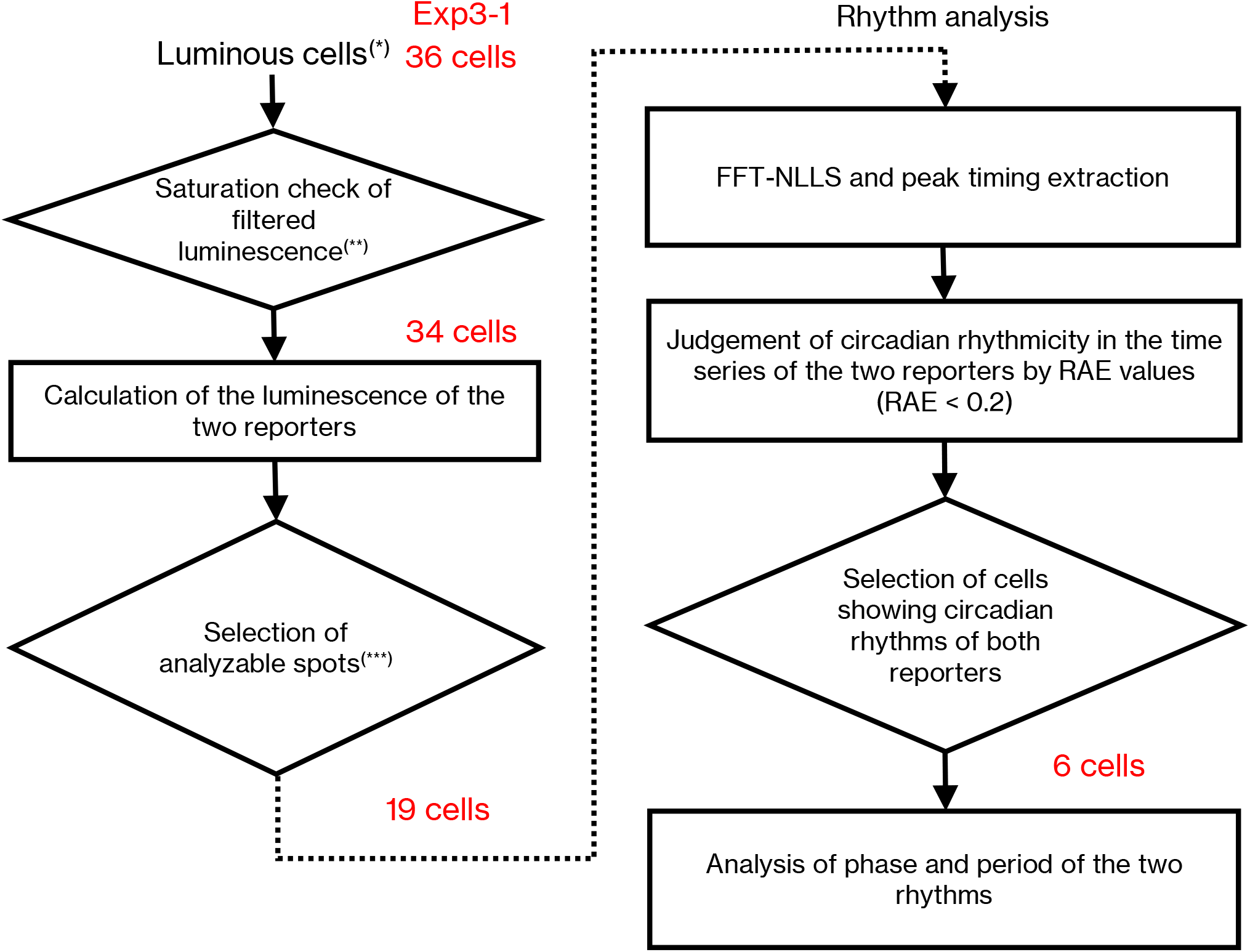
Flow diagram for the analysis of the co-transfection experiment. (*) Luminous spots without overlap are counted. (**) Luminous spots that include saturated pixels in the 6 × 6 spot area at two successive time points in the monitoring period (24–144 h) are removed. (***) Those spots with the following three criteria are selected as analyzable spots for both reporters. (1) Median of *MA*[*L_LUC+_*] > 50 photons/s and median of *MA*[*L_PtRLUC_*] > 500 photons/s. (2) Median of *MA*[*L_LUC+_*] < 5000 photons/s. (3) The number of *R*(*i*) in the range 1/20 < *R*(*i*) < 1 is more than half of the total number of datapoints. The definitions of each symbol in the criteria are as follows: *L_LUC+_* and *L_PtRLUC_* at time *i* are defined as *L_LUC+_*(*i*), *L_PtRLUC_*(*i*). The 24 h moving average of *X* at time *i* [*MA*(*X*, *i*)] is defined as follows:

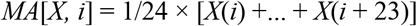 The dataset of the time series *MA*[*X*, *i*] is described as *MA*[*X*]. The luminescence ratio, *R*(*i*), of LUC+ and PtRLUC at time *i* is defined as follows.

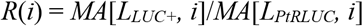

**Figure S10.**
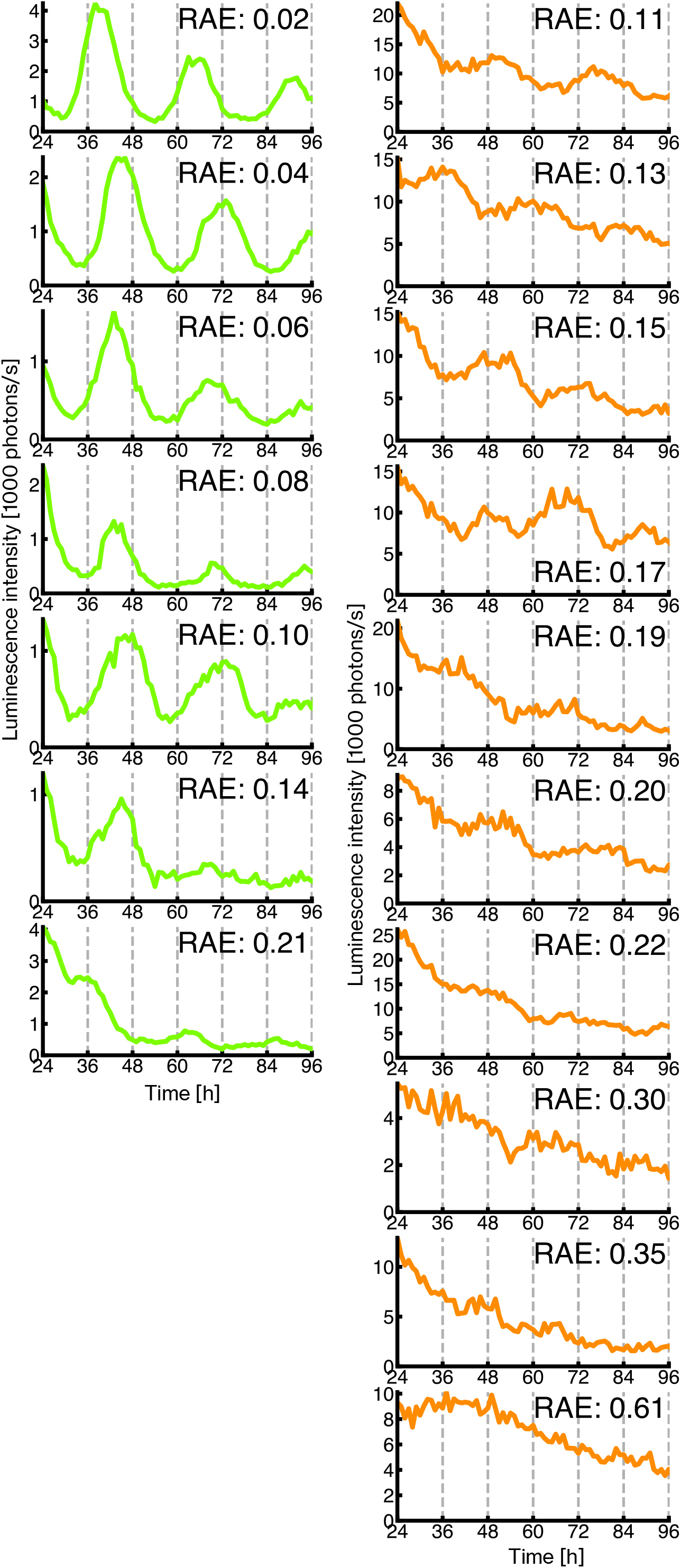
Examples of bioluminescence traces with various RAE values in single-reporter transfection experiments. Reconstructed time series of seven cells for LUC+ (light green, Exp1-1) and 10 cells for PtRLUC (orange, Exp2-1) are sorted by their RAE values. The RAE value is represented in each graph.

**Figure S11.**
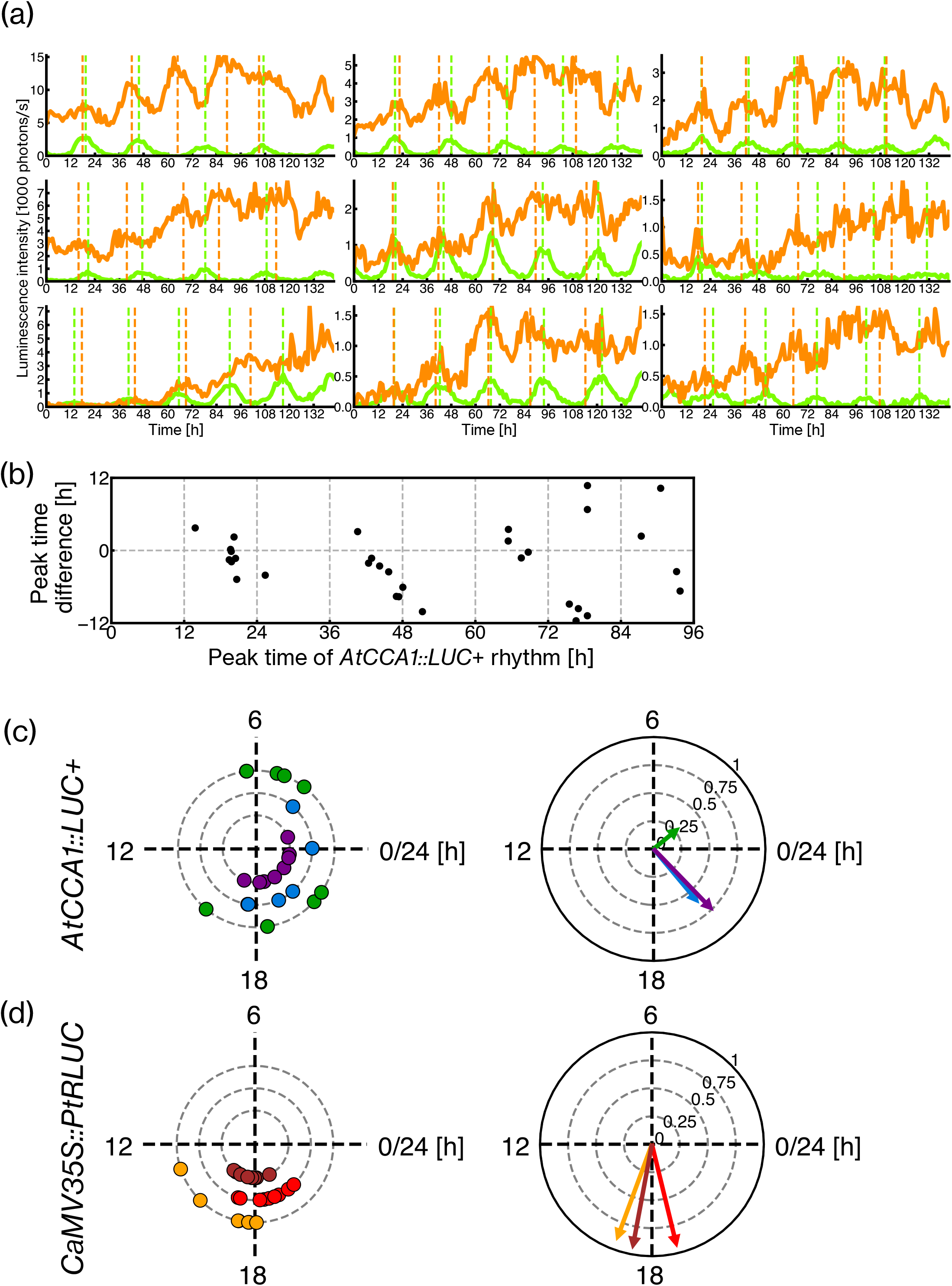
Characterization of the phase relationship of cellular bioluminescence rhythms of *AtCCA1::LUC*+ and *CaMV35S::PtRLUC* in the co-transfection experiment, Exp3-2. (a) Reconstructed time series of LUC+ (light-green line) and PtRLUC (orange line) for the nine cells showing the circadian rhythms of both reporters in Exp3-2. The dashed vertical lines show the peak times of LUC+ (light green) and PtRLUC (orange). (b) Peak time differences between *CaMV35S::PtRLUC*- and *AtCCA1::LUC*+ rhythms in Exp3-2. Data display and plotting are the same as in Figure 5a except for the range of the x-axis. Peaks that occurred in the range from 0 to 96 h are plotted because the circadian rhythmicity of *CaMV35S::PtRLUC* of most cells were lost after the range. (c, d) Plots for the peak times and their averages/synchronicity of cellular bioluminescence rhythms in Exp3-2. The peak times of the nine cells are plotted for *AtCCA1::LUC*+ (c) and *CaMV35S::PtRLUC* (d) on the left. Data display and plotting are the same as in Figure 5b,c. The peak time(s) shown in day 2 (24–48 h) is plotted on the innermost circle and those in day 3 (48–72 h) and day 4 (72–96 h) are plotted on the second and third circles, respectively.

**Figure S12.**
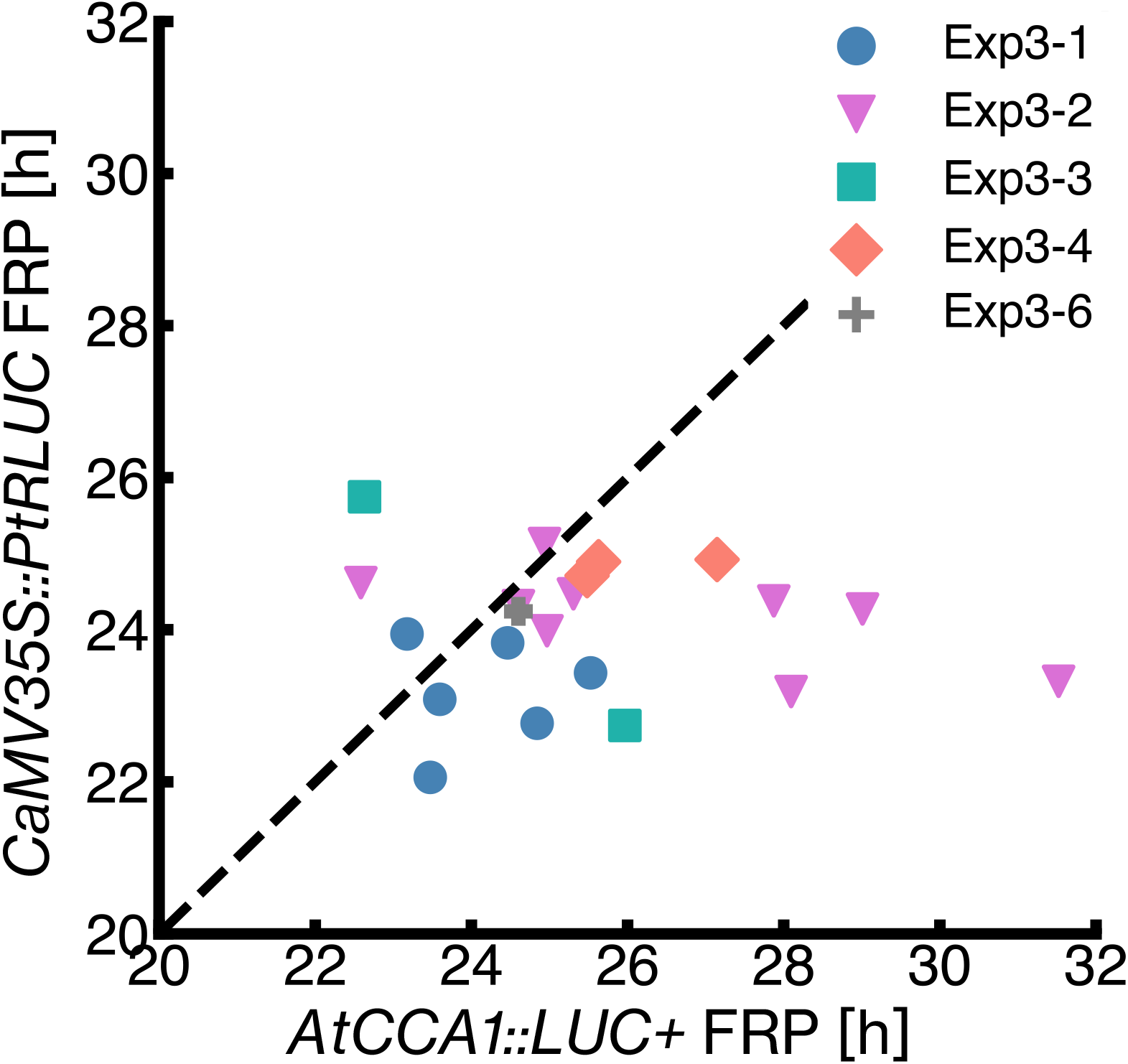
A scatterplot of the free-running period (FRP) between *AtCCA1::LUC*+ rhythm and *CaMV35S::PtRLUC* rhythm in co-transfection experiments. FRPs of the 21 cells showing the circadian rhythm of both reporters in the five co-transfection experiments (Table S1) are plotted. The same symbol (with the same color) represents cells in the same experiment. The dashed line represents where FRPs are the same between both rhythms.

**Table S1.**
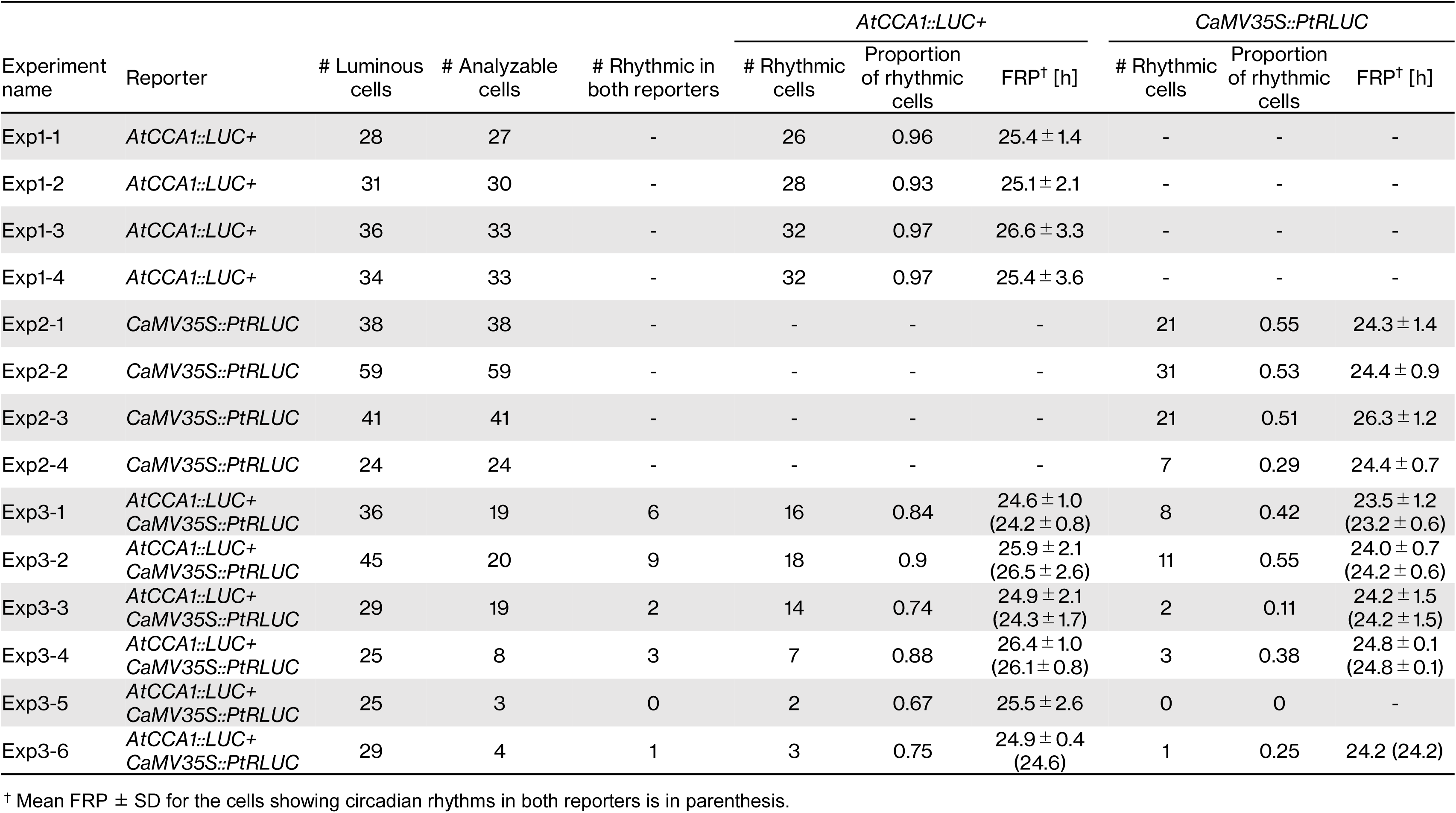
Summary of the quantitative analysis of cellular bioluminescence rhythms.

